# Time-deterministic cryo-optical microscopy

**DOI:** 10.1101/2023.08.01.551103

**Authors:** M. Yamanaka, K. Tsuji, Y. Kumamoto, S. Tamura, W. Miyamura, T. Kubo, K. Mizushima, K. Kono, H. Hirano, K. Sugiura, S. Fukushima, T. Kunimoto, K. Nishida, K. Mochizuki, Y. Harada, N. I. Smith, R. Heintzmann, T. Nagai, H. Tanaka, K. Fujita

## Abstract

We developed a technique to rapidly freeze biological systems during observation under an optical microscope, which allows for detailed optical observation at a precise time in the biological dynamics at high spatial resolution and quantifiability under low-temperature conditions. Our study further found that ion distributions and molecular conformations can be fixed in space and time, demonstrated by visualization of calcium waves rapid-frozen at an instant using fluorescent ion indicators. These results show that time-deterministic cryo-optical microscopy can suspend cellular dynamics while maintaining molecular and ionic states, allowing us to view cellular dynamics in a way that has not been possible before.

**One-Sentence Summary:** Rapid freezing of dynamic biological systems during optical observation presents time information to the fixed samples

## Main Text

Cryogenic fixation has been utilized to fix the motion of samples and their chemical state so that the sample can be placed in an environment suitable for microscopic and nanoscopic observations such as optical and electron microscopy (*1–4*). Especially in optical microscopy, the increase in the optical stability of sample molecules under cryogenic conditions allows us to improve the imaging capability in terms of quantitative detection in correlative observations and spatial resolution, providing more detailed information via multimodal approaches (*5–11*). Compared to chemical fixation, cryofixation offers the distinct advantage to preserve cellular structures (*12–14*). This preservation is essential for analysis of cellular details using super resolution fluorescence imaging techniques (*11,14–21*). In addition, cryofixation can fix the chemical state and environment in samples, such as the redox state of molecules, pH, and ion concentration, which cannot be realized by traditional chemical fixation techniques, such as using paraformaldehyde (*22–24*).

The above advantages of cryofixation have not been fully utilized in fluorescence imaging because the main purpose of cryofixation thus far has been to maintain the morphology of the samples for electron microscopy (*3*). In contrast, fluorescence microscopy observes the dynamics of biological events, where molecular or ion signaling, and the interaction of molecules and proteins are visualized together with the cell morphology and chemical status (*25–27*). Biological studies require control of the chemical and biological environments and the dynamic behaviors of samples under various stimuli, which is difficult to realize with current cryofixation techniques, such as plunge freezing and high-pressure freezing techniques (*12, 13, 28–30*).

In this study, we developed a technique to rapidly freeze living samples during observation using optical microscopy. This technique allows us to visualize the path of biological dynamics by tracing temporal information before fixation and enables us to arrest the morphological and chemical conditions of a cell at an arbitrary time during a biological event for subsequent observation using multimodal optical techniques, such as super-resolution microscopy and Raman microscopy, with enhanced chemical stability of target molecules. In addition, chemical environments such as ion distributions, pH, and redox conditions can also be fixed, allowing a more detailed investigation of chemical responses that govern biological phenomena. This technique can even fix the state of molecules, including both endogenous and exogenous molecules, such as fluorescent indicators, providing snapshots of molecular behavior at a moment of interest. Although rapid freezing techniques of a sample on a microscope stage have recently been proposed (*31–34*), immobilization of the molecular and morphological dynamics of a sample in a time-deterministic manner, which is the target in this research, is essential to allow detailed microscopic observations. Recording the sample behaviors up to the moment of fixation is a major difference from conventional fixation methods, which enables us to utilize the detailed molecular information given by fixed samples to understand dynamic cellular functions.

### On-stage chamber for rapid freezing

We developed a sample-freezing chamber that can be placed on an inverted optical microscope, as shown in Fig.1A (Text S1). A liquid cryogen composed of propane and isopentane was introduced into the sample in the chamber so that the living cells were frozen by heat exchange between the cryogen and sample (Text S2). To suppress ice crystal formation, the amount of buffer solution around the cells was reduced to a level that did not affect cell activity, and cryogen was manually applied over the sample. Using the freezing chamber, the cooling speed of 10500 °C/s was achieved, above the cooling speed for vitrification of water in a cell (10000 °C/s) (*12, 35,36*), over a temperature range between 15 °C and −100 °C (Fig. 1B and Text S3a). At this cooling speed of 10500 °C/s, the sample motion at 37 °C is presumed to be halted within around 4.5-5.0 msec (the typical ice nucleation temperature in animal cells: around −10 to −15 °C (*37*)). The cryogen used for fixation was also used to maintain the temperature of the sample under cryogenic conditions during optical observations.

**Fig. 1.**
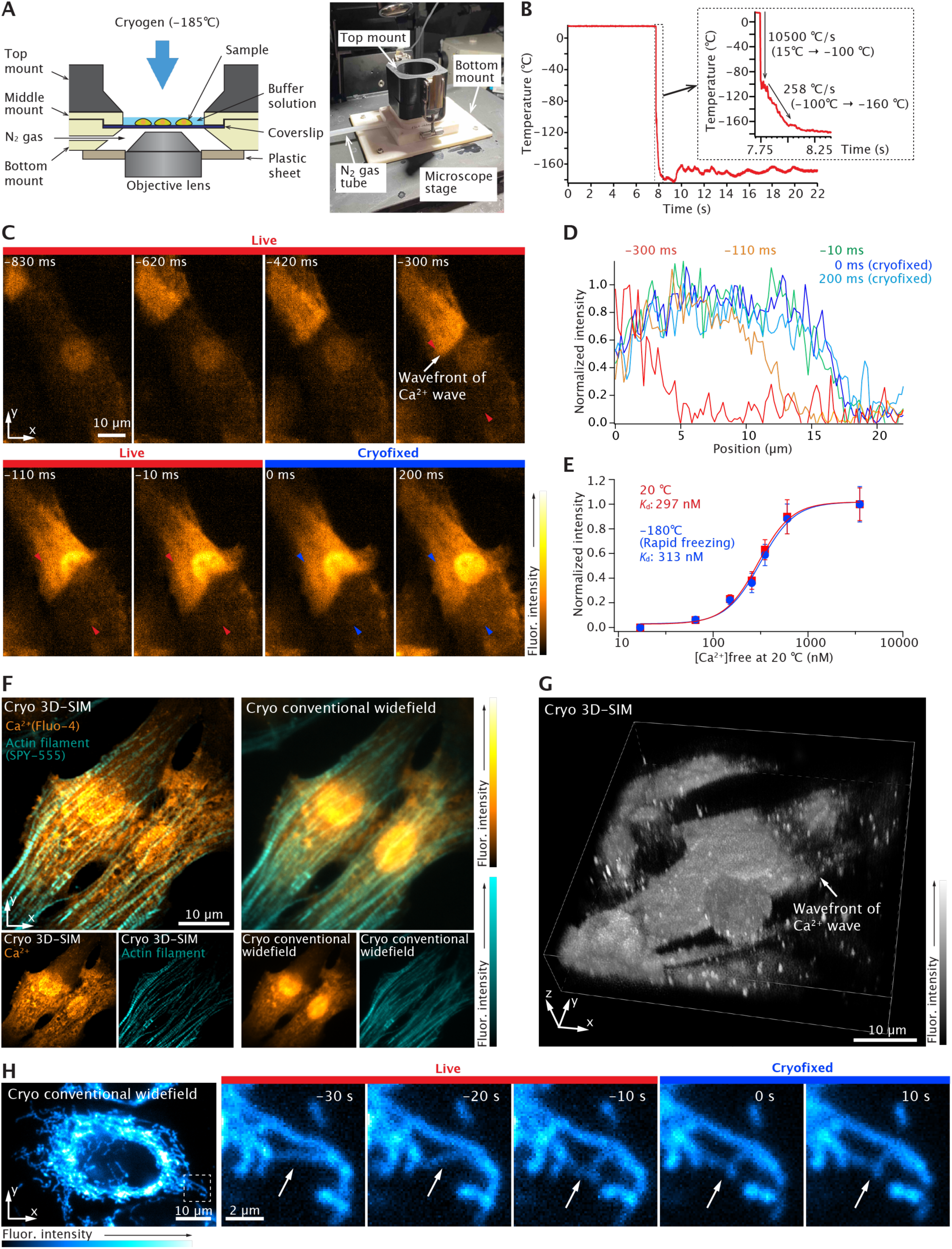
On-stage freezing chamber, cryofixation of cellular dynamics under microscopic observation, and cryogenic super-resolution imaging. (**A**) Cross-sectional schematics and photograph of on-stage freezing chamber. (**B**) The cooling speed when approximately 40-μm-thick pure water on a coverslip was frozen by a liquid cryogen (−185 °C, a mixture of liquid propane and isopentane). (**C**) Cryofixation of Ca^2+^ wave propagating in the neonatal rat cardiomyocyte (a frame rate: 100 frame/s, Movie S1). Ca^2+^ wave motion stopped at the frame of 0 ms by cryofixation. (**D**) Fluorescence intensity profiles of the line indicated with the red and blue arrows in (C). These profiles show that the leading edge of the Ca^2+^ wave moves from left to right in this plot and then stops by cryofixation. (**E**) Ca^2+^ titration curve of Fluo-4 under 20 °C and −180 °C (rapid freezing). This data was measured by using Fluo-4 in a Ca^2+^ calibration buffer (Text S4b). (**F**) Cryogenic dual-color super-resolution and conventional widefield fluorescence images of Ca^2+^ distribution and actin filaments in the neonatal rat cardiomyocytes. (**G**) Cryogenic 3D super-resolution fluorescence image of Ca^2+^ distribution in the neonatal rat cardiomyocyte (Movie S2 and S3). 3D visualization was produced by using alpha rendering (Nikon, NIS-Elements). (**H**) Cryogenic conventional widefield fluorescence images of a HeLa cell expressing DsRed in mitochondria (a frame rate: 10 frame/s, Movie S4).

### Cryofixation of cellular dynamics

Cryofixation was performed using a freezing chamber under a microscope to observe the cellular activities. Fig.1C and 1D show a series of fluorescence images of neonatal rat cardiomyocytes loaded with a calcium-ion (Ca^2+^) indicator, Fluo-4, and fluorescence intensity profiles indicating the wavefront of the Ca^2+^ wave propagating in neonatal rat cardiomyocytes (Text S3b and Text S4a). During the observation of the Ca^2+^ wave, the cryogen was introduced, and the wave motion was stopped (Movie S1). This result shows that rapid freezing can preserve not only the cell morphology and Ca^2+^ distribution, but also the chemical state of the fluorescent indicator because the contrast of the image remains largely unaltered before and after fixation. Interestingly, the dissociation constant K_d_ of Fluo-4 is known to be temperature-dependent (*38*). However, we found that the K_d_ value after rapid freezing showed the same K_d_ as before the freezing (20 °C) (Fig.1E, Text S3c and S4b). This indicates that if the freezing process is fast enough, molecular responses and spatio-temporal conditions are halted without passing through a thermal equilibrium state. During this experiment, the same optical system was used at room and cryogenic temperatures because the excitation and fluorescence spectra of Fluo-4 were not significantly different under these temperature conditions (Fig. S1).

The immobilization of the sample on the microscope stage can be fully exploited by various imaging techniques. Fig. 1F shows a super-resolution fluorescence image obtained by three-dimensional (3D) structured illumination microscopy (3D-SIM) of Fluo-4 and SPY-555, an actin probe for live cells in neonatal rat cardiomyocytes, demonstrating the super-resolution imaging of ions and associated cell behaviors that cannot be performed using conventional chemical fixation techniques (*39*) (Fig. S2, Text S3d and S4a). In fact, our result shows that the sarcomere lengths were slightly longer in the high Ca^2+^ concentration region compared to the low Ca^2+^ concentration region (Fig.S3). Ca^2+^ wave propagation in neonatal rat cardiomyocytes was halted by cryofixation during observation with a conventional widefield fluorescence microscope (Text S3d and S4a, Movie S2), and then 3D super-resolution imaging of the Ca^2+^ distribution in neonatal rat cardiomyocytes was performed using 3D-SIM (Fig. 1G, Text S3d and S4a, Movie S3). We also confirmed that the cryofixation technique froze the motions of various subcellular structures, such as the mitochondria (Fig. 1H, Text S3e and S4c, Movie S4) and lysosomes (Text S4c and Movie S5).

### Time-deterministic cryofixation

As shown above, the rapid cryofixation on the microscope stage allows us to precisely determine the time of fixation during observation, which provides temporal information on the biological event to within the duration of cryofixation (around 4.5-5.0 ms). Time-deterministic features are especially useful when combined with a technique that triggers biological events. We therefore used a caged Ca^2+^ compound that can modulate the Ca^2+^ concentration in cells by light irradiation to trigger Ca^2+^ waves in neonatal rat cardiomyocytes before cryofixation (*40, 41*) (Text S3f). As shown in Fig. 2A, Ca^2+^ propagation in the cell was frozen at 430 ms after applying the trigger pulse (Movie S6). This technique can be combined with various external stimuli, such as electrical stimulation, chemical condition changes, and mechanical stimulation (*42–44*).

**Fig. 2.**
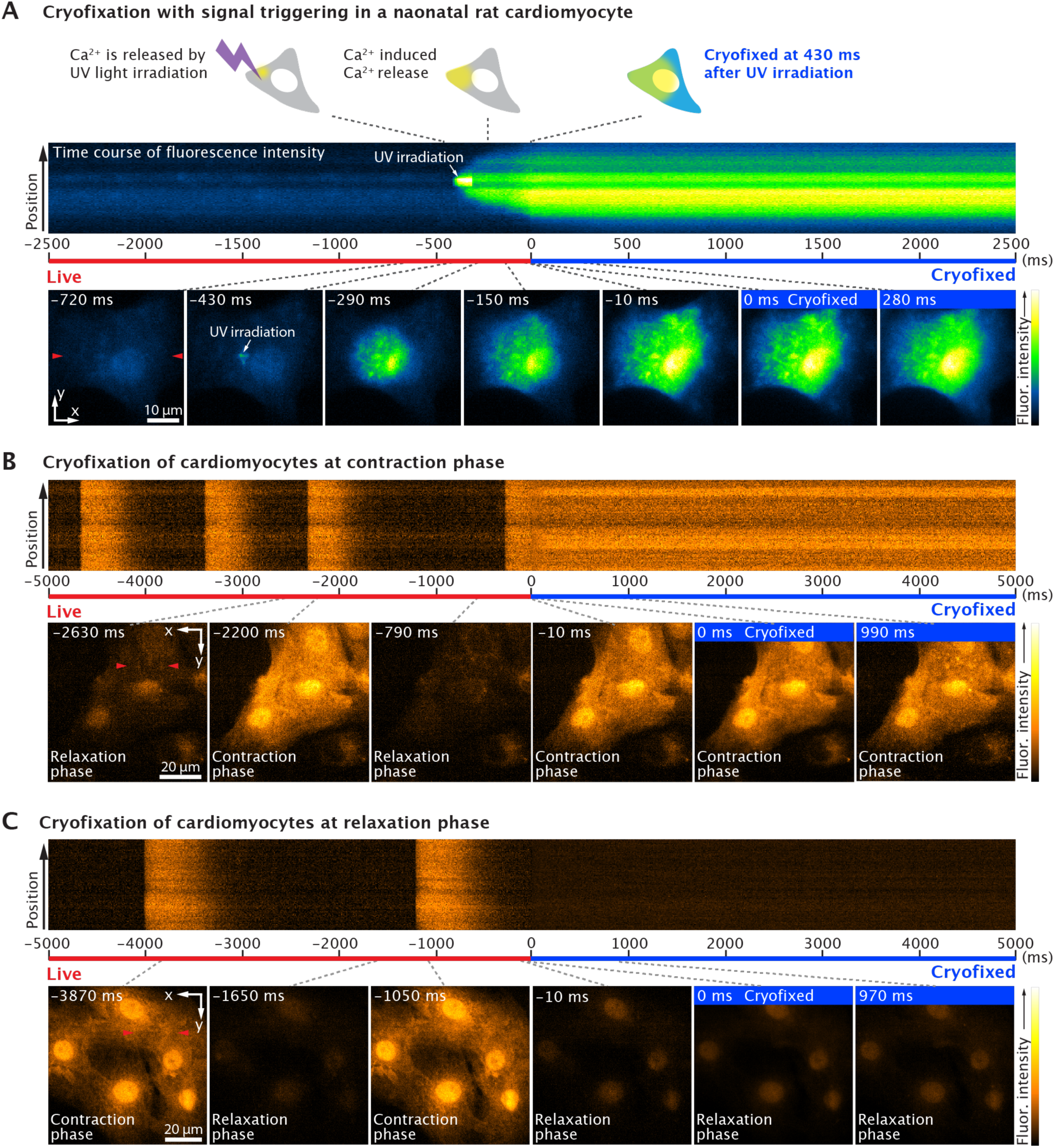
Time-deterministic cryofixation of Ca^2+^ dynamics. (**A**) Schematics of the time course of Ca^2+^ signal triggering and cryofixation, and the time course of the change of fluorescence intensity of Fluo-4 at the line indicated by the red arrows in the frame of the X-Y image at −720 ms, and X-Y images at different times. Ca^2+^ propagation stopped at 430 ms after UV light irradiation by cryofixation (Movie S6). (**B** and **C**) Cryofixation of the neonatal rat cardiomyocytes loaded with Fluo-4 during contraction and relaxation phases (Movie S7 and S8). The fluorescence images were acquired at a frame rate of 100 frame/s in all data.

As further examples, Fig. 2B and 2C show that we can selectively apply the cryogen at contraction or relaxation phases in neonatal rat cardiomyocytes, allowing the possibility to arrest heartbeat motion triggered by the cells (Text S3b, Movies S7 and S8). Neonatal rat cardiomyocytes were loaded with Fluo-4, and an increase in the free Ca^2+^ concentration in the cytoplasm during the contraction of the heartbeat motion, known as a Ca^2+^ transient, was observed. As shown in Fig. 2B and 2C, neonatal rat cardiomyocytes were frozen at the rising (contraction phase) and falling time points of fluorescence intensity (relaxation phase).

### Improved quantitative measurement

The enhanced chemical stability of the molecules under cryogenic conditions is beneficial for quantitative measurements (*8, 9*). Under ambient conditions, it is difficult to quantify dynamic samples. However, this is much easier under cryogenic conditions. As demonstrated in Fig. 3A, long-time exposure of a cryofixed sample provided a higher signal-to-noise ratio (SNR) in fluorescence imaging compared to the limited exposure time available in dynamic imaging (Text S3g). High resistance to photobleaching under cryogenic conditions (Fig. S4) was also beneficial for improving the SNR. The improvement in the SNR allowed us to enhance the quantification capability of the experiment. For example, we confirmed the SNR, which can provide 0.2 nM accuracy in Ca^2+^ measurement, using Fluo-4 with 10 s exposure under −170 °C, demonstrating a 17-fold improvement compared to measurement conditions that could be used in live cell imaging (33 ms here, at 20 °C) (Fig. S5).

**Fig. 3.**
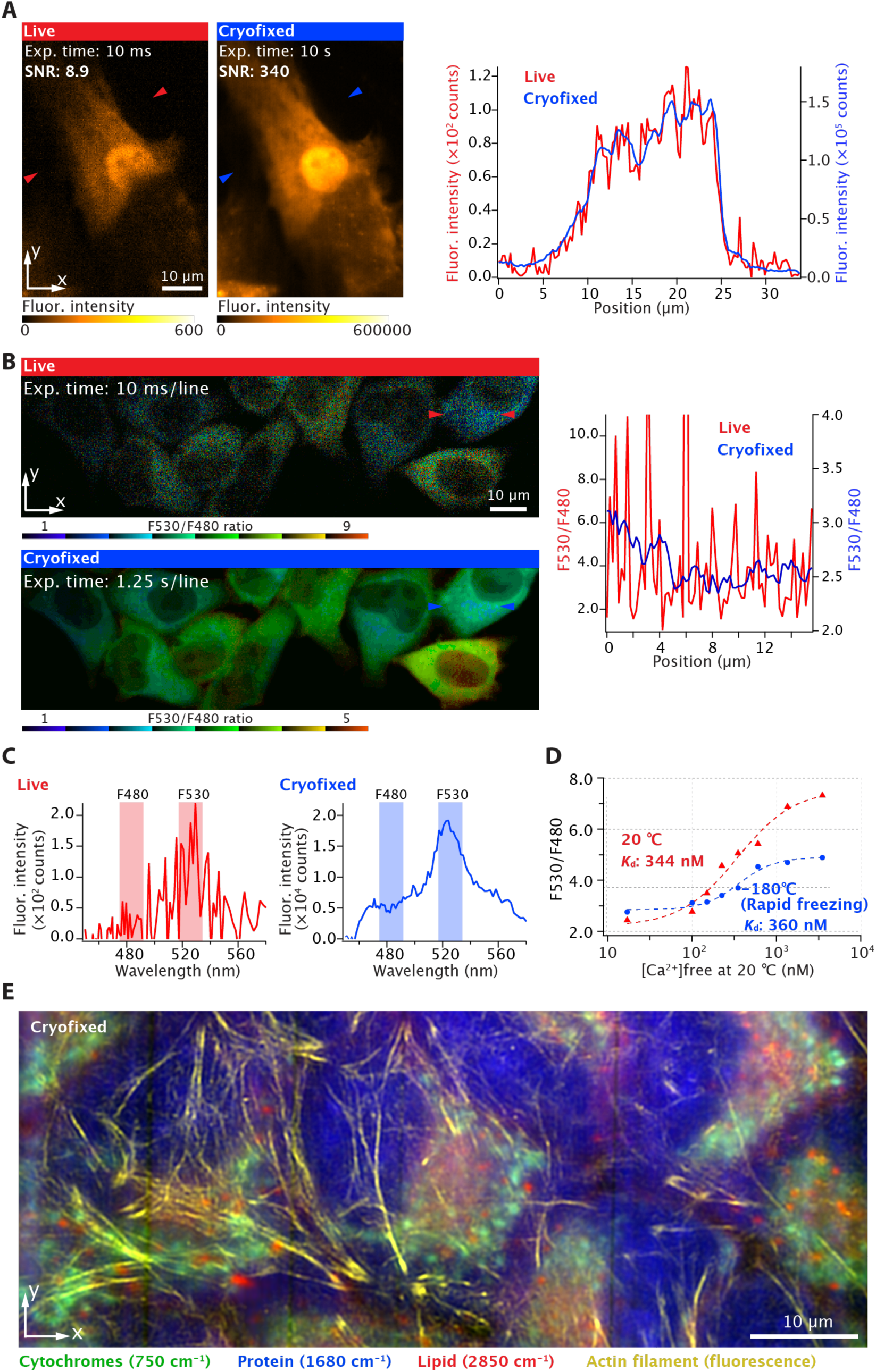
Image quantification ability is improved by ultra-rapid fixation and cryogenic imaging. (**A**) Fluorescence images of Fluo-4 labeled neonatal rat cardiomyocyte and fluorescence intensity line profiles of the lines indicated by the red and blue arrows in the fluorescence images. By increasing the exposure time by a factor of 1000, the SNR was improved from 8.9 to 340. The experimental data are the same as that shown in Fig. 1C. The fluorescence image of the cryofixed sample with the exposure equivalent to 10 s was generated by integrating 1000 fluorescence images with an exposure time of 10 ms under cryogenic conditions. (**B**) Ratiometric fluorescence images of HeLa cells expressing YC3.60 acquired with a slit-scanning hyperspectral fluorescence microscope before and after cryofixation. Here, we utilized MetaMorph software (Molecular Devices) to generate the ratiometric image using 8 shades of color in a look-up table shown at the bottom of the image. To facilitate the identification of cellular morphology, the intensities of each color in the image were adjusted to correspond with fluorescence intensities observed in the fluorescence image of Venus. Fluorescence ratio line profiles of the line indicated by the red and blue arrows in the ratio images. The ratio values in the line profiles were calculated from the fluorescence intensity images of ECFP and Venus (Fig. S7). Similar to (A), the fluorescence image with the equivalent long exposure time under cryogenic conditions was generated by the integration of 125 fluorescence images with an exposure time of 10 ms/line. (**C**) Representative fluorescence spectra of YC3.60 in the sample shown in (B). The increase of the exposure time allows us to improve SNR in the spectrum measurement. (**D**) Ca^2+^ titration curve of YC3.60 before and after cryofixation, which was measured by using YC3.60 dispersed in Ca^2+^ calibration buffer solution. (**F**) Cryogenic spontaneous Raman and super-resolution fluorescence imaging of HeLa cells. The super-resolution fluorescence image was taken with 3D-SIM and shown in yellow color. Raman intensities at 750, 1680, and 2850 cm^-1^ are mapped, representing the distributions of cytochromes (green), protein (blue), and lipid (red), respectively.

We also examined a ratiometric fluorescent Ca^2+^ probe, yellow cameleon 3.60 (YC3.60), which is a pair of variants of cyan fluorescent protein, ECFP, and yellow fluorescent protein, Venus, connected to Ca^2+^-sensitive calmodulin (*45*). Förster resonance energy transfer (FRET) between ECFP and Venus is facilitated by the capture of Ca^2+^ by calmodulin, and the ratio of the fluorescence intensity of ECFP and Venus can quantitatively show the Ca^2+^ concentration in cells. Fig. 3B shows a ratiometric fluorescence image of HeLa cells expressing YC3.60 in the cytosol and fluorescence ratio profiles obtained from the lines indicated by red and blue arrows in Fig. 3B, acquired using a slit-scanning hyperspectral fluorescence microscope (*46*) (Text S3h, Fig. S6). The cells were cryofixed at 36 s after histamine stimulation. Long-time exposure under cryogenic conditions allowed us to measure the fluorescence spectrum of YC3.60 with a higher SNR, as shown in Fig. 3C, providing a high quantification capability for Ca^2+^ imaging. We also compared the K_d_ value of YC3.60 under the room temperature and the cryogenic condition at −180 °C after the rapid freezing mentioned above. As expected from the previous experiments described above, the K_d_ value did not change significantly under these conditions. However, the fluorescence ratios increased at low Ca^2+^ concentrations and, on the other hand, decreased at high Ca^2+^ concentrations after cryofixation (Fig. 3D, Text S3i).

Although further investigations are needed to clarify the mechanism behind the change in fluorescence ratio after rapid freezing, this might be explained by changes in physicochemical properties, such as the increase in FRET efficiency due to the elongated fluorescence lifetime under low temperatures, a larger increase in the quantum yield of ECFP compared to that of Venus, changes in the emission spectrum shapes and possibly absorption spectrum shapes (Fig. S8-S10). Despite the changes in these physicochemical properties, ratiometric imaging at a high SNR for the quantitation of Ca^2+^ concentration was achieved, indicating that various FRET-based probes can also be used with time-deterministic cryo-optical microscopy. The fluorescence ratio in cryogenic cell imaging was lower than those shown in the Ca^2+^ titration curve, which we attribute to the difference in the surrounding environmental conditions of YC3.60 in HeLa cells and the Ca^2+^ calibration buffer solution.

### Multimodal imaging capability with a different temporal resolution

Time-deterministic cryofixation is also useful for multimodal studies because switching between different modalities does not cause a time discrepancy between modes and allows observation of the same precise moment even using modalities with different temporal resolutions, such as fluorescence and Raman microscopy (*47, 48*) (Fig. S11). Here, we demonstrated this by multimodal imaging of the same state of HeLa cells using fluorescence 3D-SIM and slit-scanning Raman microscopy (*47*) (Fig. 3E, Fig. S12, Text S3j). Actin filaments in HeLa cells were labeled with SPY555 in the same manner, as shown in Fig. 1F. After cryofixation, a super-resolution fluorescence image of actin filaments and high-resolution Raman images of cytochrome, protein, and lipid distributions were acquired. The image acquisition times for fluorescence 3D-SIM and slit-scanning Raman microscopy were 0.75 s and 25 min, respectively. Although only two microscopic imaging modalities were used in this study, a wide variety of optical microscopic modalities, such as autofluorescence and coherent Raman imaging techniques, can be added, allowing us to increase multiplexing of observation targets or add further orthogonal imaging data (*49, 50*).

## Discussion

We demonstrated rapid cryofixation of living cells under observation by optical microscopy, in which the morphological and chemical dynamics of the cells were fixed at precisely defined time. The fixation speed can reach a level at which ice crystal formation in a cell is prevented during freezing (vitrification). The freezing procedure is similar to that used in plunge-freezing techniques, where the amount of liquid at the sample surface is reduced before introducing the sample into the cryogen for vitrification with rapid freezing. Through our study, we also confirmed that rapid freezing overall preserved cellular shapes, although slight changes in morphology could be measured, and these effects were found to be mitigated by adding the cryoprotectant trehalose to the buffer solution (Fig. S13). The change in cellular shapes is presumably due to the difference in volume change of the cell body and the extracellular solution during cooling. The fact that the addition of trehalose reduced the shape change suggests that the formation of ice crystal far from the observation area or minute ice crystals in the extracellular solution, which are invisible in the optical observation, could pressure the sample. Note that, in our experiments, we did not observe sample destruction even without cryoprotectant (Fig. S13).

Fast freezing allows us to precisely time observation moments within the dynamical processes of biological events from different aspects, which can lead to new discoveries and improved understanding of biological phenomena. Furthermore, the rapid freezing allows preserving native structures in soft and non-membranous organelles, e.g. as formed by liquid-liquid phase separation (*14, 51, 52*). It is also possible to combine other imaging and analytical modalities, such as electron microscopy and mass spectroscopy, to provide more comprehensive insights into biological phenomena. Additionally, fluorescence labeling methods commonly used for chemically fixed samples, such as immunostaining and *in situ* hybridization, can be applied after freeze substitution of cryofixed cells (*53*). Compared with chemical fixation, cryofixation produces fewer morphological and chemical artifacts (*12–14*). Moreover, time-deterministic cryofixation allows more tools to be combined than previously possible, providing an accurate and comprehensive snapshot of biological events. The accurate and detailed data could also be useful as the ground truth for machine learning approaches (*54–56*).

One of the remarkable findings of this study is that the K_d_ value of fluorescent Ca^2+^ probes is maintained after rapid freezing, which indicates that the molecular-level interactions in the cell can be frozen without passing through a thermal equilibrium state. Therefore, our results suggest that rapid freezing can stop chemical reactions and can be effective for the selection of a precise immobilization timepoint. This technique may allow us to analyze the moment of chemical reactions, which can be applied to study chemical dynamics for many different purposes. As the fixation of Ca^2+^ distribution in 3D was successfully performed, this technique can also be extended to fix the activities of other ions, including those for membrane potential and signaling molecules. This technique can be used to fix states of the cell that relate to otherwise intangible information such as pH, redox state, and temperature, which can be detected using functional fluorescent probes (*57–59*).

Optical measurement under cryogenic conditions achieved a high SNR in optical detection. Especially in the imaging of chemically unfixable targets, such as ions or other signaling messengers, more details of their intracellular redistribution can be revealed without being limited by short exposure times needed for traditional time-resolved imaging. This advantage is also significantly beneficial for enhancing the imaging performance of super resolution fluorescence microscopy (*39, 60–64*) which can benefit greatly from relaxing constraints on acquisition times. The concept and design of fluorescent probes can also be altered when considering the usage and optimization of their performances under cryogenic condition. For the same reason, this technique is also useful for improving the detection sensitivities of optical techniques that require the detection of low-light signals, such as Raman microscopy. Cryogenic Raman imaging using Raman tags can be used to observe low concentrations of small molecules in live cells that are difficult to observe using fluorescence labeling (*65–67*). SNR and detection sensitivities could be further improved if an immersion objective lens with a high numerical aperture is used, which allows us to improve the spatial resolution (*10, 68–70*). Moreover, the implementation of adaptive optics can further optimize the spatial resolution and signal levels (*71*).

In the past, several benefits of cryogenic optical microscopy have been discussed and demonstrated; however, these have not been available to the wider imaging communities because of complexity and cost for cryofixation and cryo-observation (*4–21, 28–34*). The technique developed in this study enables practical usage of cryogenic optical microscopy and realistically extends optical imaging by providing these advantages demonstrated here to all research fields relying on optical microscopy.

## Acknowledgments

The authors thank M. Hayakawa for the isolation of cardiomyocytes from neonatal rats, S. Kawano for the development of SIM software, and S. Sakakihara for the preparation of SIM masks.

## Author contributions

Conceptualization: K.F.

Methodology: K.F., M.Y., Y.K., Y.H., H. T., T.N.

Investigation: M.Y., K.T., W.M., S.T., T.Kubo, K.Mizushima, K.K., H.H., K.S., S.F., T.Kunimoto, K.Mochizuki, K.N., N.I.S., K.F.

Visualization: M.Y., K.T., W.M., T.Kubo, K.Mizushima

Software: T.Kubo, R.H., K.T.

Funding acquisition: K.F., K.T.

Project administration: K.F., M.Y.

Supervision: K.F.

Writing – original draft: K.F., M.Y., K.T., S.T., K.Mizushima

Writing – review & editing: N.I.S., T.N., Y.H., H.T., R.H.

## Data and materials availability

All data used in this paper are available upon reasonable request.

## List of Supplementary Materials

Supplementary Text

Fig. S1 to S13

Movie S1 to S8

References (72-83)

