## Supplementary Information for "Time-deterministic cryo-optical microscopy"

**The PDF file includes:**

Supplementary Text
Figs. S1 to S13
References (72-83)

**Other Supplementary Materials for this manuscript include the following:**

Movies S1 to S8

**Table of Contents**

**Supplementary Text**

**S1.** On-stage freezing chamber

**S2.** Liquid cryogen

**S3.** Experimental details and conditions in Fig. 1 to 3 in the main text

a) Fig. 1B

b) Fig. 1C, 1D, 2B, and 2C

c) Fig. 1E

d) Fig. 1F and 1G

e) Fig. 1H

f) Fig. 2A

g) Fig. 3A

h) Fig. 3B and 3C

i) Fig. 3D

j) Fig. 3E

**S4.** Sample preparations and staining

a) Neonatal rat cardiomyocyte

b) Fluo-4 and YC3.60 in calcium calibration buffer solution

c) HeLa cell

d) Protein preparation of recombinant YC3.60

**Fig. S1** Excitation and fluorescence spectra of Fluo-4 in HEPES buffer

**Fig. S2** Dual color 3D structured illumination microscope (3D-SIM)

**Fig. S3** Sarcomere lengths in a cryofixed neonatal rat cardiomyocyte

**Fig. S4** Reduction of photobleaching effects of Fluo-4 under cryogenic condition

**Fig. S5** Improvement of signal-to-noise ratio by increasing an exposure time under cryogenic condition

**Fig. S6** Hyperspectral slit-scanning fluorescence microscope

**Fig. S7** Fluorescence images of HeLa cells expressing YC3.60 before and after cryofixation

**Fig. S8** Fluorescence lifetime images of HeLa cells expressing YC3.60 at room temperature and cryogenic conditions

**Fig. S9** Fluorescence images of HeLa cells expressing ECFP and Venus before and after

cryofixation
**Fig. S10** Fluorescence spectra of YC3.60, ECFP, and Venus before and after cryofixation
**Fig. S11** Multimodal SIM/Raman microscope
**Fig. S12** Raman spectra of a HeLa cell under cryogenic conditions
**Fig. S13** Cryogenic fluorescence images of HeLa cells with and without cryoprotectant
**Movie S1** Cryofixation of $Ca^{2+}$ waves in neonatal rat cardiomyocytes (Fig. 1C).
**Movie S2** Cryofixation of $Ca^{2+}$ waves in neonatal rat cardiomyocytes without cryoprotectant (Fig. 1G)
**Movie S3** 3D visualization of cryofixed $Ca^{2+}$ waves in neonatal rat cardiomyocytes (Fig. 1G)
**Movie S4** Cryofixation of mitochondria in HeLa cells without cryoprotectant (Fig. 1H)
**Movie S5** Cryofixation of lysosomes in HeLa cells without cryoprotectant
**Movie S6** Time-deterministic cryofixation of $Ca^{2+}$ waves induced by uncaging $Ca^{2+}$ from a caged $Ca^{2+}$ compound with UV light irradiation (Fig. 2A).
**Movie S7** Time-deterministic cryofixation of neonatal rat cardiomyocytes at the contraction phase (Fig. 2B)
**Movie S8** Time-deterministic cryofixation of neonatal rat cardiomyocytes at the relaxation phase (Fig. 2C)
**References**

### Supplementary Notes

#### S1. On-stage freezing chamber

The sample freezing chamber consisted of three parts: top, middle, and bottom mounts, as shown in Fig. 1(A). A sample coverslip was placed in the indented area of the bottom mount and held by pressing it down from above. The top mount was equipped with two hinges. After a sample is mounted, it can then be fixed by applying liquid cryogen from the upper side, and the sample freezing chamber allows the storage of liquid cryogen during observation. The area between the sample coverslip and the objective lens was filled with nitrogen gas, which was kept in a chamber with a plastic sheet and an objective-lens-sized hole in the center to prevent frost condensation due to the low temperature. This simple design facilitates easy handling of this sample freezing chamber, significantly reducing the time required to exchange a sample to approximately 30-60 s by using spare mounts. This sample freezing chamber can be attached to any type of standard inverted microscope stage using appropriate adapters.

The top, middle, and bottom mounts were fabricated using commercially available 3D printers. The top mount was made of polylactic acid, which does not break at -196 °C and has low thermal conductivity (0.13 W/m·K). The upper mount was designed as a hollow structure with 90% air (0.0257 W/m·K when 20 °C) to improve the thermal insulation of the top mount. The top mount was fabricated using a fused-filament fabrication-type 3D printer (UltiMaker, Ultimaker S5). PLA (Ultimaker, Tough PLA) was used for the top mount. The middle and bottom mounts are in direct contact with a sample coverslip; therefore, the surface in contact with the sample coverslip must be flat and smooth to prevent the buffer solution of the sample from leaking from the mount. For this reason, these mounts were fabricated using a stereolithography-type 3D printer (Formlabs, Form3), which enables more precise and smoother fabrication than fused filament fabrication-type 3D printers. Rigid 10K resin (Formlabs) was used as a stereolithography-type 3D printer to produce the middle and bottom mounts.

A sample (cells on a 10×10 mm, 18×18 mm, or 25 mm diameter quartz coverslip (thickness of 0.17 ± 0.02 mm)) was removed from the cell culture medium in a cell culture dish, and the liquid on the bottom side of the coverslip was carefully wiped off with paper such as lens cleaning paper with ethanol. Immediately thereafter, the sample was mounted in an on-stage freezing chamber, and a buffer solution, such as Hank's balanced salt solution (HBSS) or phosphatebuffered saline (PBS), was added to prevent the sample from drying out while it was being prepared for measurement.

The experimental data shown in this paper were obtained using a Nikon Ti2-E microscope. After completing the measurement preparation and activating the axial autofocus system of the Ti2-E microscope, the buffer solution was aspirated using a pipette. Under an optical microscope, liquid cryogen was manually applied to freeze a sample at any given time, such as when $Ca^{2+}$ waves or $Ca^{2+}$ transients occur.

### S2. Liquid cryogen

For our experiments, a mixture of liquid isopentane and propane (temperature: about -185 °C) was mainly used. Only for multimodal imaging (Fig. 3E), liquid propane (temperature: about -185 °C) was used.

Liquid cryogens were prepared using a semiautomated custom-built cryogen preparation apparatus (Cryovac, CV-Z-LTMIX100). To prepare the mixture of liquid isopentane and propane, liquid isopentane was poured into a glass beaker and placed in the preparation apparatus. The liquid isopentane in the beaker was cooled with liquid nitrogen ($LN_2$), and propane gas was introduced into the cooled liquid isopentane and liquefied in the liquid isopentane. Throughout the liquefaction process of propane gas, the mixture was continuously stirred. When the preparation finished, the beaker was taken out of the apparatus, and the mixture was cooled down to -185 °C with $LN_2$ and kept until needed for the experiments.

To prepare liquid propane, an empty glass beaker was placed in the apparatus and cooled with $LN_2$. Propane gas was blown onto the inner surface of the beaker for liquefaction. Then, the liquid propane was cooled down to -185 °C with $LN_2$ in the same way as in the case of a mixture of liquid isopentane and propane.

### S3. Experimental details and conditions of data in Fig. 1 to 3 in the main text

#### a) Fig. 1B

To measure a cooling rate when a liquid cryogen is applied to a sample, a 25 μm diameter type-K thermocouple (ANBE SMT Co., KFT-25-100-100) was placed in approximately 40-μm-thick pure water on a coverslip in an on-stage freezing chamber. The thermocouple was connected to a thermocouple amplifier electrical circuit (Analog Devices, AD8495) with a theoretical cutoff frequency of 8 kHz and the signals were recorded with our homemade software at 1 kHz sampling rate. The temperature was measured by applying a liquid mixture of isopentane and propane (temperature: -185 °C) onto the thermometer and pure water (temperature: 15 °C). In the result, the temperature dropped below 0 °C within 1 ms, and the cooling rate from 15 °C to -100 °C was 10500 °C/s, which is higher than that required for vitrification of water (10000 °C/s) (*12, 35,36*). At a temperature below -100 °C, the cooling rate became dramatically decreased (258 °C/s). This is presumably due to the presence of excessive volume of water on the thermocouple, and the cooling rate would be maintained if water volume could be controlled to be less than the current one. When the liquid cryogen was applied directly onto the thermocouple, we confirmed that the cooling rate of 10000 °C/s was maintained at a temperature down to the vitrification point of water (-135 °C). Although ice crystals may form in a sample, it is considered that their sizes were much smaller than the spatial resolution of optical microscopes used in this study. This conclusion is supported by the absence of no apparent damage resulting from the formation of ice crystals, as observed in the experimental results presented in this paper.

**b) Fig. 1C, 1D, 2B, and 2C**

The Fluo-4 loaded neonatal rat cardiomyocytes were rapidly frozen under a microscopic observation with a conventional inverted widefield fluorescence microscope (Nikon, Ti2-E) equipped with a mercury lamp. The $Ca^{2+}$ wave propagation stopped at the image frame at 0 ms, as shown in Fig. 1C and 1D, and it was considered that the liquid cryogen contacted the sample at the image. The contact timing of liquid cryogen on the sample was determined by finding the image frame in the time-series images where the fluorescence intensity suddenly increased over the entire area of the image. This is because the fluorescence intensities of fluorescent dyes are known to increase at low temperatures owing to an increase in their quantum yields (*72, 73*). In this case, the fluorescence intensity of Fluo-4 increased over the entire area of the image frame at 0 ms, and the enhancement factor was 1.18 compared to that in the immediate previous image frame. We also confirmed that the image contrast was preserved after cryofixation, indicating that the sample was cryofixed while preserving both the $Ca^{2+}$ distribution and the $K_d$ of Fluo-4. In the same manner, $Ca^{2+}$ transients in Fluo-4 loaded neonatal rat cardiomyocytes were observed and the sample were frozen at the contraction and relaxation phases. Here, we also confirmed that image contrast was preserved after cryofixation. Although further investigations are needed to elucidate the details of the slight difference in the image contrast before and after cryofixation, this may be attributed to contamination of cellular autofluorescence, which is also enhanced under cryogenic condition, and the difference of physiochemical properties between Fluo-4 and cellular molecules under cryogenic condition. If this is the case, employing spectral unmixing techniques or a calcium indicator with longer excitation and emission wavelengths would mitigate this issue significantly.

For visualization, the fluorescence intensity of each image was normalized to the fluorescence intensity of the selected area in the observed cell. As the fluorescence intensity was enhanced under cryogenic conditions, we applied normalization for the fluorescence images separately before and after cryofixation. First, the background signal values before and after cryofixation were obtained from an area where there were no cardiomyocytes in each image, and then subtracted from each image. To normalize the fluorescence images before cryofixation, we chose one bright area in the fluorescence image of an observed cardiomyocyte just before cryofixation and took the mean intensity value. Fluorescence images before cryofixation were divided by the mean intensity value. To normalize the fluorescence images under cryogenic conditions, mean intensity values at the same position in each image as that before cryofixation were used. In contrast to normalization of the images before cryofixation, each fluorescence image under cryogenic condition was divided by the mean intensity values taken in each fluorescence image. This was performed to compensate for the change in fluorescence intensity caused by temperature fluctuation or increase after cryofixation. We noticed that the sample position drifted slightly after cryofixation, probably because of temperature changes. Therefore, we compensated for the drift motion by estimating the displacement from the cross-correlation of the phase components in the Fourier transformation of the images (*74*). The excitation and detection wavelengths were 464-499 nm (Semrock, FF01-482/35-25) and 516-556 nm (Semrock, FF01-536/40-25), respectively. The sample was observed with an NA0.7 dry objective lens (Nikon, CFI Plan Fluor 60XC), and fluorescence images were acquired using an sCMOS camera (Hamamatsu Photonics, ORCA Flash4.0 V3). The autofocus system of the Nikon microscope was activated during the observations; the exposure time was 10 ms and the frame rate was 100 frames/s. In this experiment, we added trehalose in a buffer solution as a cryoprotectant. The concentration of trehalose was 200 mM.

**c) Fig. 1E**

The dissociation constants ($K_d$) of calcium indicator Fluo-4 were measured under 20 °C (room temperature) and -180 ℃ (cryofixed). In the measurements under 20 °C (room temperature) and -180 ℃ (cryofixed), we repeated the following procedure: 1) we measured fluorescence signals of a Fluo-4 solution with a certain $Ca^{2+}$ concentration under 20 °C, 2) froze it rapidly with a liquid cryogen and measured fluorescence signals under -180 ℃, and then 3) changed the Fluo-4 solution to one with different $Ca^{2+}$ concentration. The $K_d$ values were obtained by fitting the measured data with a sigmoidal function (*75, 76*). It was confirmed that a $K_d$ value similar to that in Ref. 38 of the main text was obtained at the room temperature and the shape of the $Ca^{2+}$ titration curve was preserved after cryofixation. This result supports the preservation of the image contrast of the Fluo-4 loaded neonatal rat cardiomyocytes after cryofixation, as demonstrated in Fig. 1C. Fluorescence signals were measured with a conventional widefield inverted fluorescence microscope (Nikon, Ti2-E) equipped with a mercury lamp. Fluorescence signals were collected with an NA0.45 dry objective lens (Nikon, S Plan Fluor ELWD 20x) and acquired with a spectrophotometer (Princeton Instruments, Acton SP2500) equipped with an electron multiplying charged-couple device (EMCCD) camera (Andor, iXon Ultra 888). The excitation and detection wavelength bands were 464-499 nm (Semrock, FF01-482/35-25) and 516-556 nm (Semrock, FF01-536/40-25), respectively.

**d) Fig. 1F and 1G**

Images were obtained by 3D-SIM under cryogenic conditions (Fig. S2). Because the actin probe SPY555 allowed us to label F-actin, the gaps between actin filaments in the sarcomeres of neonatal rat cardiomyocytes were also more clearly observed using 3D-SIM (*39*). In this observation, the exposure time was 25 ms for structured illumination at each angle and phase, and 15 raw images with three angles and five phases per angle were acquired. Trehalose was added to the buffer solution as a cryoprotectant for the experiment in Fig. 1F, whereas trehalose was not added to the buffer solution for the experiment in Fig. 1G.

In Fig. 1G, $Ca^{2+}$ wave propagation in neonatal rat cardiomyocytes was stopped by cryofixation under a microscopic observation with a conventional inverted widefield fluorescence microscope (Nikon, Ti2-E) equipped with a mercury lamp, and then performed 3D imaging with 3D-SIM. To acquire a 3D image, the autofocus system of the Nikon microscope was activated during observation, and axial scanning was performed by changing the Z-direction offset position of the autofocus system in steps using homemade software. The step size along the Z direction was 285 nm. The other experimental parameters were identical to those shown in Fig. 1F. In Movie S2, the fluorescence intensities of each image were normalized for visualization in the same manner as shown in Fig. 1C.

Image acquisition and reconstruction of SIM images were performed using homemade software in MATLAB. This software allows us to perform pre-processing before reconstructing a SIM image from raw images. The pre-processing process involves normalizing fluorescence intensities among raw images and correcting small sample drifts through phase correlation. For SIM image reconstruction, a pre-processed image set is spatially Fourier transformed, and the high spatial frequency components are separated and reassigned in the spatial frequency space through a matrix-based unmixing process. If there is a mismatch between an experimental result and a theoretical estimate of reconstruction parameters, the mismatch produces artifacts in a reconstructed image. To reduce the artifacts, this software also allows us to optimize the reconstruction parameters, such as the phases and frequencies of structured illumination (*77*). A Wiener filter and weighting averaging were also applied to each frequency component to reduce the Poisson noise in the frequency space and enhance the high-frequency components achieved by structured illumination (*78*). Finally, a super-resolution image was reconstructed by inverse Fourier transforming the resultant frequency distributions.

**e) Fig. 1H**

HeLa cells expressing DsRed in mitochondria were observed with a conventional inverted widefield fluorescence microscope equipped with a mercury lamp. As shown in Fig. 1H, the mitochondria moved under the room temperature and stopped by cryofixation. Although the cell shape was slightly deformed by cryofixation, it was not significant as shown in the enlarged views in Fig. 1H. This result indicated that, if multiple targets are observed by using different fluorescent probes for live cell imaging, it would be possible to analyze the situation, location, and distribution of multiple organelles at the certain time point of biological events with improved SNR. The excitation and detection wavelength bands were 527-552 nm (Nikon, excitation filter of TRITC filter cube set) and 577-632 nm (Nikon, emission filter of TRITC filter cube set), respectively. The sample was observed with an NA0.7 dry objective lens (Nikon, CFI Plan Fluor 60XC) and fluorescence images were acquired with an sCMOS camera (Hamamatsu Photonics, ORCA

Flash4.0 V3). The exposure time was 100 ms. The fluorescence intensities of each image were normalized for visualization purpose in the same manner as that in Fig. 1C. Trehalose was not added to the buffer solution in this experiment.

**f) Fig. 2A**

Neonatal rat cardiomyocytes were loaded with Fluo-4 and caged calcium compound (Tocris, DMNPE-4 AM-caged-calcium). During widefield fluorescence observations with a mercury lamp, UV light was focused on a cardiomyocyte for 100 ms to uncage $Ca^{2+}$ and then they were rapidly frozen at 430 ms after UV irradiation. At the time of the UV irradiation, the bright spot appeared in the fluorescence imaging as shown in the fluorescence image at -430 ms. After uncaging $Ca^{2+}$, the $Ca^{2+}$ wave propagated through $Ca^{2+}$ induced $Ca^{2+}$ release (CICR), and then the $Ca^{2+}$ wave was stopped during the propagation by cryofixation. In this experiment, we added trehalose in a buffer solution as a cryoprotectant. The sample drift was compensated, and the fluorescence intensities of each image were normalized for visualization purposes in the same manner as that in Fig. 1C.

A nanosecond pulsed laser system with a center wavelength of 355 nm (Spectra-Physics, EONE-355-1YHY-W) was used as the light source for the optical stimulus. The laser beam was introduced into an acousto-optic modulator (AOM) (IntraAction, ASM-802B8), used as a high-speed shutter, and the 1st order diffracted beam was introduced into a conventional widefield Nikon Ti2-E inverted fluorescence microscope. The opening and closing of the AOM shutter were controlled by applying square waveform electrical signals from the function generator to the AOM driver. UV light and excitation light from a mercury lamp for widefield fluorescence imaging were combined with a dichroic mirror and subsequently introduced into the microscope. UV light was focused on the sample with an NA0.7 dry objective lens (Nikon, CFI Plan Fluor 60XC) with approximately 50% transmittance at a wavelength of 355 nm. Fluorescence signals were collected with the same objective lens, and fluorescence images were recorded with an sCMOS camera (Hamamatsu Photonics, ORCA-Flash4.0 V3). The excitation and detection wavelength bands for the widefield fluorescence observations were 464-499 nm (Semrock, FF01-482/35-25) and 516-556 nm (Semrock, FF01-536/40-25), respectively.

**g) Fig. 3A**

Using the data shown in Fig. 1C, we evaluated the improvement in the SNR under cryogenic conditions. As described in the figure caption, to generate a fluorescence image with an exposure time equivalent to 10 s, we integrated 1000 fluorescence images with an exposure time of 10 ms under cryogenic conditions, and then calculated the SNR under the assumption that noise follows a Poisson distribution. Although we evaluated the SNR of the fluorescence image with an exposure time equivalent to 10 s here, it is feasible to extend the measurement time under cryogenic conditions for the further improvement of SNR.

**h) Fig. 3B and 3C**

HeLa cells expressing YC3.60 were observed using a hyperspectral slit-scanning fluorescence microscope with line illumination (Fig. S6) (*46*). During hyperspectral imaging, HeLa cells were stimulated with histamine and then frozen rapidly, capturing the intracellular free $Ca^{2+}$ concentration change occurring upon histamine stimulation. From the time course of fluorescence intensity in the stimulated HeLa cells, we confirmed that HeLa cells were frozen when the free $Ca^{2+}$ concentration was increased by histamine stimulation. Although the fluorescence ratio decreased after cryofixation, the line profiles of the fluorescence ratio shown in Fig. 3B indicate that the spatial variation of the fluorescence signal was preserved after cryofixation. Trehalose was not added to the buffer solution in this experiment.

HeLa cells were immersed in HBSS buffer before histamine stimulation, and then HBSS buffer containing 10 μM histamine was added for histamine stimulation. After gentle pipetting of the HBSS buffer containing histamine and a subsequent waiting time of 30 s, the HBSS buffer was removed, and the sample was frozen rapidly. The excitation intensity was 6.8 μW on the sample plane and the exposure time per line was set to 10 ms to achieve an image acquisition time of 4 s in this experiment, which is relatively slow, but still allows us to observe the oscillation of fluorescence intensities induced by histamine stimulation.

Prior to image reconstruction from the measured fluorescence spectra of YC3.60, we measured the baseline offset value (bias level) of the EMCCD camera with its shutter closed, and then subtracted it from the measured fluorescence spectra. Fluorescence images were reconstructed using the averaged fluorescence signals at the wavelength bands where ECFP and Venus had their fluorescence peaks. The spectral bands chosen for the fluorescence images of ECFP and Venus were 475-492 nm and 517-534 nm, respectively. To reduce the remaining background signals, the average intensity values of regions without HeLa cells were obtained in each reconstructed intensity image, and the estimated background signals were then subtracted from each reconstructed intensity image. Then, YFP/CFP ratio images were generated using MetaMorph software (Molecular Devices).

**i) Fig. 3D**

In the same way as for Fig. 1E, fluorescence spectra of YC3.60 were acquired with a spectrophotometer equipped with an EMCCD camera (Andor, iXon Ultra 888) under different $Ca^{2+}$ concentrations. To calculate the fluorescence ratio, we used the same wavelength regions as those used for reconstructing the fluorescence ration images in Fig. 3B. The excitation wavelength band was 400-410 nm (Semrock, FF01-405/10-25). The fluorescence signals were detected in a wavelength region above 461 nm (Semrock, LP03-458RU-25). An NA0.7 dry objective lens (Nikon, CFI Plan Fluor 60XC) was used for this observation.

**j) Fig. 3E**

HeLa cells labeled with SPY-555-actin (Spirochrome) were observed with the developed microscope (Fig. S11). The sample was mounted on a custom-made cryostage with a cooling/heating function (Linkam Scientific) and rapidly frozen with liquid propane (-185 °C). After rapid freezing, the sample temperature was controlled with the cryostage, and the liquid propane (boiling point: -42 °C) was removed from the sample by vaporization under the temperature setting of the cryostage at -40 °C. This is because it was not possible to detect Raman signals from HeLa cells under the existence of propane, which provides significantly stronger Raman signals than HeLa cells. Although we used a mixture of liquid isopentane and propane for rapid freezing in the other experiments, propane alone was used in this experiment due to the requirement of vaporization after freezing (the boiling point of isopentane at 28 °C would be too high to vaporize). Trehalose was not added to the buffer solution in this experiment.

In this multimodal imaging, fluorescence 3D-SIM and spontaneous Raman images were acquired sequentially. For fluorescence 3D-SIM imaging, the sample was illuminated with a laser light at a wavelength of 561 nm (intensity: 35.8 $kW/cm^2$). Fifteen raw images were obtained at three different illumination angles and five different phases per angle. The total image acquisition time was 750 ms. After 3D-SIM imaging, the same 561-nm excitation light was illuminated on the sample for 675 s to photobleach the fluorescent probe and reduce the strong and broad fluorescence backgrounds in Raman imaging. Raman signals of the HeLa cells were detected under line illumination with laser light at a wavelength of 532 nm. The excitation intensity was 300 kW/cm$^2$ and the exposure time was 10 s/line. Raman images were reconstructed after removing the noise and bias background components by applying singular value decomposition (SVD) to the data obtained (*47*).

### S4. Sample preparations and staining

#### a) Neonatal rat cardiomyocyte

All animal experiments described in this study were conducted in accordance with the *Guide for the Care and Use of Laboratory Animals* (8th edition, National Academies Press, Washington DC, 2011) following the approval of the Animal Research Committee at the Kyoto Prefectural University of Medicine (approval No: M2022-238). Primary cultures of neonatal rat cardiac myocytes were prepared as previously described with some modifications (*79, 80*). Briefly, 2- or 3-day-old Wistar rats (Japan SLC, Inc.) were anesthetized using intraperitoneal injection with 0.1 mg/kg of medetomidine, 3.0 mg/kg of midazolam, and 5.0 mg/kg of butorphanol, and the hearts were removed and placed in $Ca^{2+}$- and $Mg^{2+}$-free PBS. The aorta was discarded, and the atria and ventricles were minced under aseptic conditions. The small pieces were enzymatically digested three times for 10 min each with 10 mL of PBS containing 0.2% type II collagenase (Worthington) at 37 °C. Cell suspensions from each digestion were pooled, centrifuged at 800 rpm for 5 min, and resuspended in Dulbecco's modified Eagle's medium (DMEM) (FUJIFILM Wako Pure Chemical) supplemented with 10% fetal bovine serum (FBS) (Nichirei biosciences inc.) and 1% PSG antibiotic mix (100 U/mL penicillin, 100 μg/mL streptomycin, 2 mM l-glutamine) (FUJIFILM Wako). To eliminate non-myocyte cells from the preparation, cells were pre-plated twice in a 100-mm culture dish for 45 min at 37 °C in a humidified atmosphere containing 5% $CO_2$-95% room air. After the pre-plating step, the cell suspension containing myocytes was collected and plated on gelatin-coated coverslips in DMEM containing 10% FBS and 1% PSG. Twenty-four hours after plating, cells were washed with fresh DMEM for experiments.

The $Ca^{2+}$ indicator Fluo-4 AM (AAT Bioquest, 20550 or Chemical Dojin, 342-90961) was used for $Ca^{2+}$ imaging, as shown in Fig.1C, 1F, 1G, 2, and 3A. To prepare the staining solution, solid Fluo-4 AM was dissolved in dimethyl sulfoxide (DMSO), and its concentration was adjusted to 1 mM. The Fluo-4 AM solution was mixed with nonionic detergent Pluonic® F-127 (Biotium,

59004), and then diluted to 1 μM with HBSS (FUJIFILM Wako Pure Chemical, 082-08961) to prepare a staining solution. Neonatal rat cardiomyocytes were washed three times with HBSS, immersed in Fluo-4 staining solution for 15 min, and washed three times with HBSS.

The live-cell F-actin probe SPY-555-actin (Spirochrome) was used to image actin filaments (Fig. 1F). Solid SPY-555 was dissolved in 50 μL of DMSO to prepare a stock solution, and then the stock solution was diluted 1000-fold with the cell culture medium to prepare a staining solution. Neonatal rat cardiomyocytes were immersed in the staining solution for 1 hour and washed three times with HBSS.

To perform $Ca^{2+}$ imaging with a caged calcium compound (Tocris, DMNPE-4 AM-caged calcium), both the caged calcium compound and Fluo-4 were loaded into neonatal rat cardiomyocytes. The caged calcium compound was dissolved in DMSO at a concentration of 100 mM. The caged calcium compound solution was mixed with a mixture of Fluo-4 and the nonionic detergent Pluonic® F-127 (Biotium, 59004). The mixture was diluted with HBSS to prepare the staining solution. The concentrations of Fluo-4 and caged calcium compounds in the staining solutions were 1 and 10 μM, respectively. Neonatal rat cardiomyocytes were stained in the same manner as for Fluo-4 staining.

**b) Fluo-4 and YC3.60 in calcium calibration buffer solution**

A calcium calibration buffer solution kit (Thermo Fisher Scientific, C3008MP) was used to prepare free $Ca^{2+}$ buffer solutions with $Ca^{2+}$ concentrations ranging from 17 to 10000 nM. To measure $K_d$ of Fluo-4, Fluo-4 pentapotassium salt (ATT Bioquest, 20555) was dissolved in a buffer solution containing 10 mM EGTA, 100 mM KCl, and 30 mM MOPS (Thermo Fisher Scientific, C3008MP) and was mixed with 39 μM free $Ca^{2+}$ buffer solution (Thermo Fisher Scientific, C3008MP) to prepare the free $Ca^{2+}$ buffer solutions with different $Ca^{2+}$ concentrations. The concentration of Fluo-4 was 5 μM. As in the case of Fluo-4, YC3.60 was dispersed in a free $Ca^{2+}$ buffer solution for the measurement of $K_d$. Concentration of YC3.60 was 3.6 μM (Text S4d).

**c) HeLa cell**

HeLa cells were grown in the same culture medium as used for neonatal rat cardiomyocytes and cultured on coverslips under the conditions of 5% $CO_2$ and 37 °C.

To observe the mitochondria in HeLa cells (Fig. 1H, Movie S4), pcDNA3 encoding DsRed, which fused CoxVIII signal peptides, was introduced into HeLa cells, using cationic lipid-mediated transfection. For transfection, HeLa cells were grown to 50-60% of confluence in 6-well plates. Then, reduced-serum medium Opti-MEM (Thermo Fisher Scientific, 31985062) containing 1 μg fluorescent protein plasmid DNA and cationic lipid-based transfection reagents (1μL Lipofectamine 3000 reagent and 5 μL P3000 reagent (Thermo Fisher Scientific, L3000008)) were added to each well, and HeLa cells were incubated for 1 day in cell culture.

The live-cell lysosome probe LysoTracker Deep Red (Thermo Fisher Scientific, L12492) was used to observe the distribution and dynamics of lysosomes. Solid LysoTracker Red was dissolved in DMSO and diluted with cell culture medium to prepare a 75 nM staining solution. HeLa cells were immersed in the staining solution for 1 hour and washed three times with HBSS.

For multimodal SIM/Raman imaging, the HeLa cells were cultured on coverslips and stained with SPY-555-actin (spirochrome). The staining condition was the same as that described in the section on neonatal rat cardiomyocytes (Text S4a).

To perform ratiometric $Ca^{2+}$ imaging (Fig. 3B), HeLa cells stably expressing yellow cameleon 3.60 (YC3.60) in the cytosol were used.

**d) Protein preparation of recombinant YC3.60.**

To yield the recombinant protein, *E. coli* [JM109 (DE3)] was transformed using $pRSET_B$ vectors encoding YC3.60 and cultured in 200 mL liquid LB medium at 23 °C for 60-72 hours. *E. coli* was collected by centrifugation, resuspended in buffer containing 50 mM Tris-HCl (pH 8.0) and 20 mM imidazole, and crushed using a French press. The supernatant of the centrifuged bacterial crushing solution was adsorbed onto Ni-NTA resin in an open column, washed with buffer for resuspension, and eluted with buffer containing 50 mM Tris- HCl (pH 8.0) 250 mM imidazole. Finally, the buffer was replaced with 10 mM EGTA in 100 mM KCl, 30 mM MOPS (pH 7.2) using PD-10 column (Cytiva).

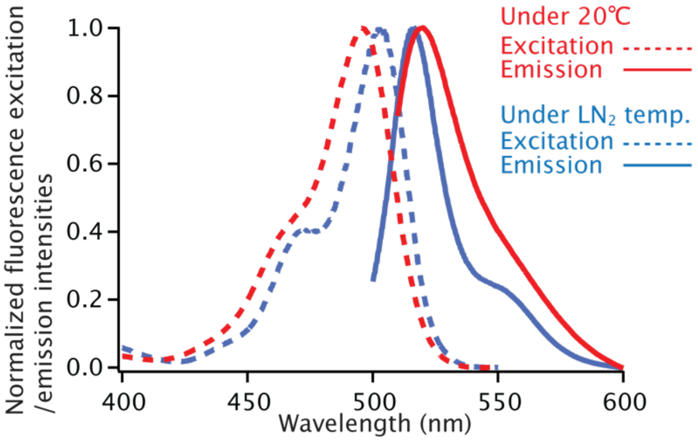


**Fig. S1.**

**Excitation and fluorescence spectra of Fluo-4 in HEPES buffer solution.** The excitation and fluorescence spectra were measured with a spectrofluorometer (Hitachi, F-7000). The measurement of those spectra under $LN_2$ temperature was performed with an optional unit for measurements at low temperature (Hitachi, 5J0-0112/4J1-0104).

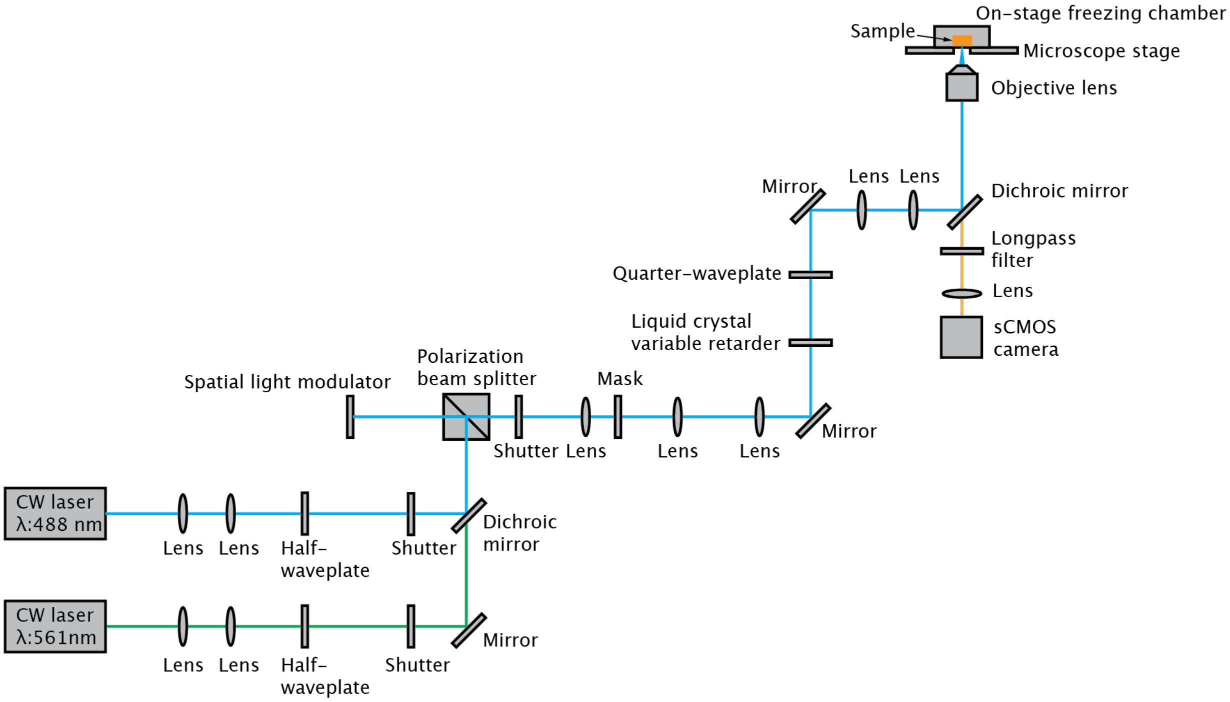


**Fig. S2.**

**Dual color 3D structured illumination microscope (3D-SIM).** The configuration of the optical setup for this microscope is similar to that previously reported in Ref. 39. The excitation wavelengths are 488 nm (Coherent, Genesis MX488-1000) and 561 nm (Spectra Physics, Excelsior 561). In this setup, a fluorescence image is acquired separately at each excitation wavelength. The polarization angle of the excitation beam is adjusted with the half-waveplate so that the excitation beam is reflected by the polarization beam splitter to a reflective spatial light modulator (SLM) (Forth dimension displays, SXGA-3DM) with 1280×1024 pixels. Periodic patterns for structured illumination displayed on the SLM produce refracted beams. The refracted excitation beam passes through the polarization beam splitter, and then only the $0^{th}$ and $\pm 1^{st}$ order refracted excitation beam passes through the mask placed at the focus position of the lens. The phases of the refracted excitation beams were adjusted to maximize the contrast of the structured illumination formed on the sample plane by using a liquid crystal variable retarder (Thorlabs, LCC1413-A) and a quarter waveplate. For this SIM, we used a Nikon Ti2-E inverted microscope equipped with the automatic axial-drift compensation system (Nikon perfect focus system). The refracted excitation beams were introduced into the microscope body and reflected to the objective by a dichroic mirror (For 488 nm excitation: Di03-R488-t3-25x36 (Semrock), 561 nm excitation: Di03-R561-t3-25x26 (Semrock)). The excitation beams were focused on the pupil plane of the objective lens. An NA0.7 dry objective lens (Nikon, CFI Plan Fluor 60XC) was used for Fig. 1F and 1G. Fluorescence signals were detected with an sCMOS camera (Hamamatsu Photonics, ORCA Flash4.0 V3) mounted on the side port of the microscope body.

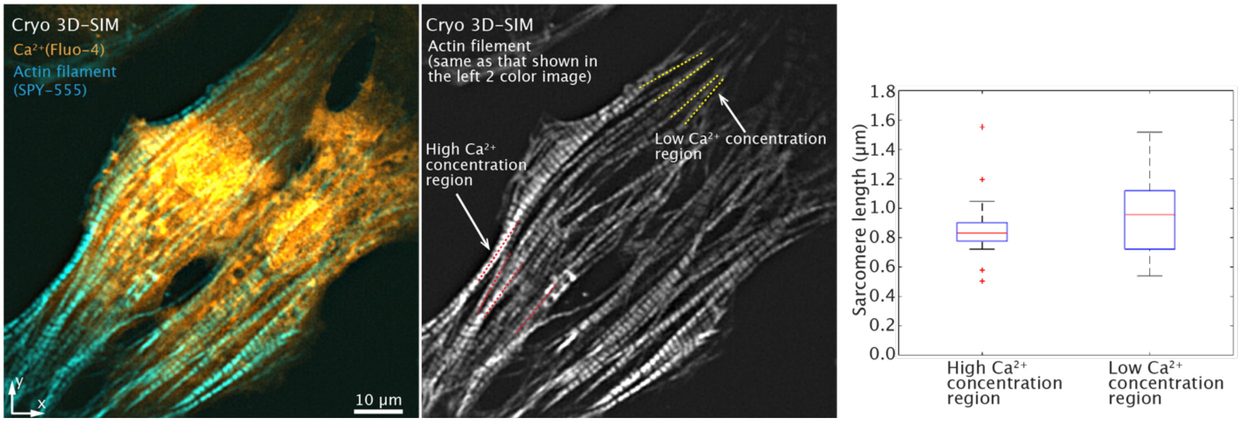


**Fig. S3.**

**Sarcomere lengths in a cryofixed neonatal rat cardiomyocyte.** We examined the sarcomere lengths in the cryofixed neonatal rat cardiomyocyte of the dual color SIM data used for Fig. 1F. The sarcomere lengths were measured in 5 actin filaments in each of the low and high $Ca^{2+}$ concentration regions, as indicated by the yellow and red dotted lines indicated in the fluorescence image of actin filaments. Our analysis showed that the sarcomere lengths were slightly longer in the high $Ca^{2+}$ concentration region compared to the low $Ca^{2+}$ concentration region (*81*). The box plot shows the distribution of sarcomere lengths in high and low $Ca^{2+}$ concentration regions and outliers are indicated by the red crosses.

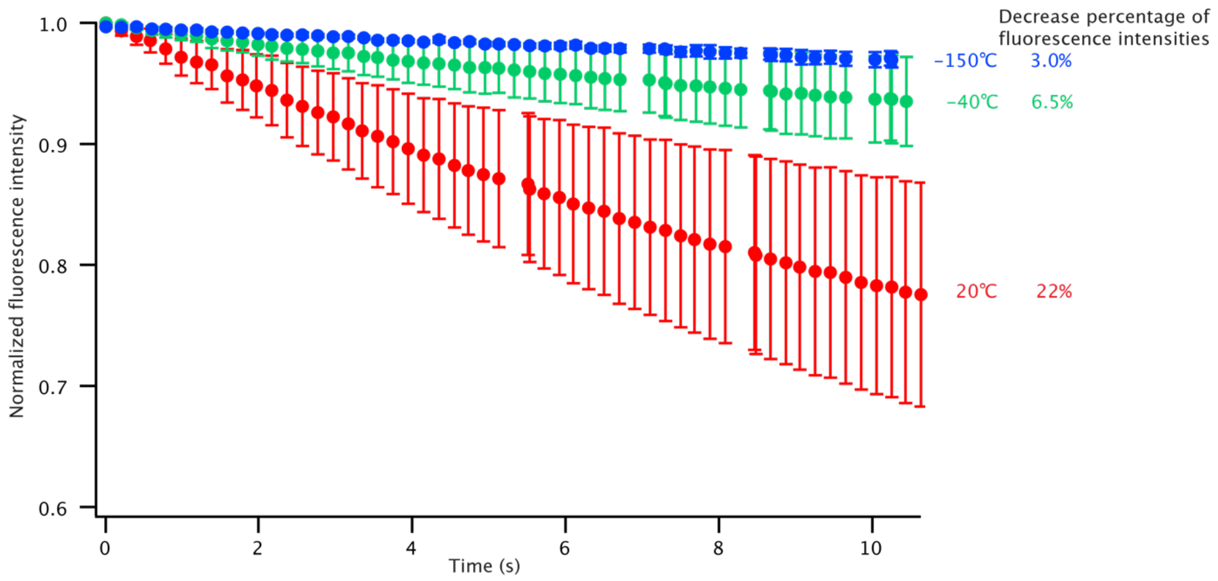


**Fig. S4.**

**Reduction of photobleaching effects of Fluo-4 under cryogenic condition.** To confirm the reduction of photobleaching effects by decreasing the temperature of a sample, Fluo-4 loaded HeLa cells at different temperatures were imaged every 200 ms for 10 s, and then the decrease of fluorescence signals was observed. In this experiment, samples were mounted in a customized cryostage (Linkam Scientific) with a temperature control function and then the samples were observed at 20, -40, and -150 °C. The cryostage was placed on the microscope stage of a conventional widefield inverted fluorescence microscope, and fluorescence images were recorded with a CCD camera. Fluo-4 was excited with a mercury lamp at the excitation intensity of 55 W /cm$^2$, which is a typical excitation intensity used for structured illumination microscopy. The excitation and detection wavelength bands were 464-499 nm (Semrock, FF01-482/35-25) and 516-556 nm (Semrock, FF01-536/40-25), respectively.

The average fluorescence intensities and standard deviations were calculated from the fluorescence images of 10 HeLa cells, and the results are plotted. During the time-course observation, we confirmed that some HeLa cells were optically stimulated, and their fluorescence signals increased (*82, 83*). For the evaluation of photobleaching, we omitted the optically stimulated HeLa cells. From the result, we confirmed that the photobleaching effect is reduced about 7 times by decreasing the temperature from 20 °C to -150 °C. This is because fluorescent molecules become more thermally stable at lower temperatures due to reduced photochemical reaction rates (*8, 9*). These results indicate that the photobleaching effect is reduced by freezing, and improved photostability helps to improve the signal-to-noise ratio by allowing the increase of exposure time for fluorescence imaging under cryogenic conditions.

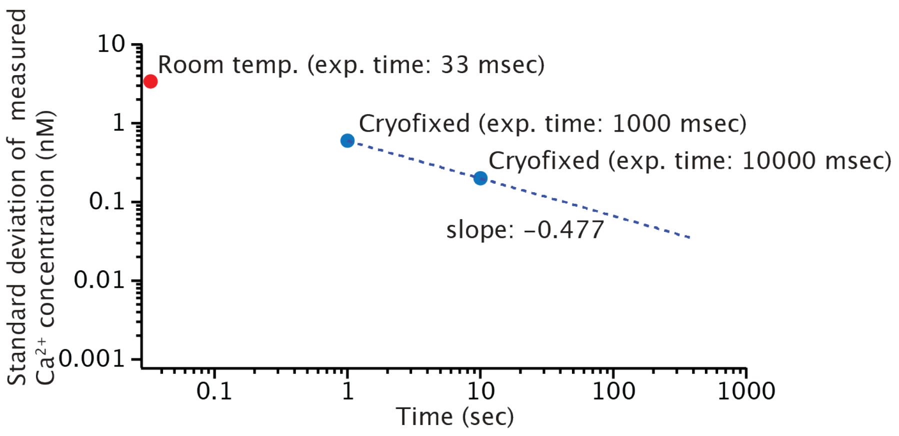


**Fig. S5.**

**Improvement of signal-to-noise ratio by increasing exposure time under cryogenic conditions.** The signal-to-noise ratio with a long exposure time under cryogenic conditions was evaluated by measuring fluorescence signals of Fluo-4 in free $Ca^{2+}$ buffer solutions with a $Ca^{2+}$ concentration of 100 nM, assuming a $Ca^{2+}$ concentration in a cell is 100 nM. The excitation laser at 488 nm was focused into the sample solution with an NA0.45 dry objective lens (Nikon, S Plan Fluor ELWD 20X). The excitation intensity was 55 μW/μm$^2$. The fluorescence signals were measured under 20 °C (room temperature) and -170 °C (cryofixed) with a slit confocal microscope equipped with a spectrophotometer and an EMCCD camera (Andor, iXon Ultra 888). From the measured fluorescence signals, standard deviations were calculated under the assumption that noises follow a Poisson distribution, and then the value of standard deviations were converted to $Ca^{2+}$ concentrations (in units of nM) by performing the following calculation: standard deviation value/signal value × $Ca^{2+}$ concentration in the sample solution (100 nM). The result indicated that the increase of the exposure time under cryogenic conditions allows us to improve the measurement accuracy of $Ca^{2+}$ concentration. As shown in this graph, the slope of the data under cryogenic conditions was -0.477, which was obtained by fitting the data with a power function. The improvement factor of the standard deviation is nearly proportional to the square root of the increase factor of the exposure time. This result indicates that the fluorescence signals of cryofixed samples were measured with almost negligible photobleaching effects.

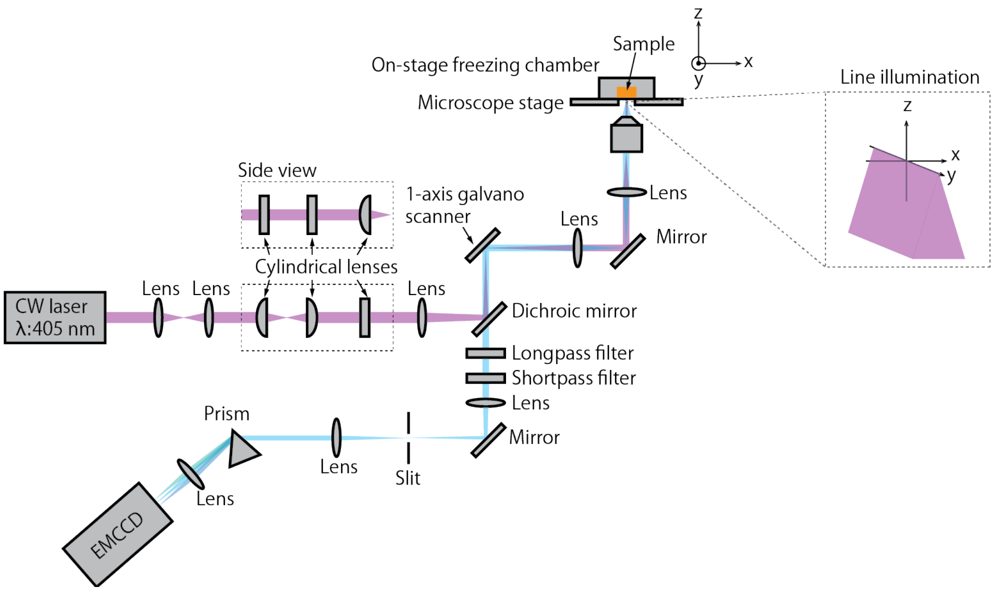


**Fig. S6.**

**Hyperspectral slit-scanning fluorescence microscope.** The optical configuration of the hyperspectral slit-scanning fluorescence microscope is similar to that described in Ref. 46. The excitation wavelength was 405 nm (HÜBNER photonics, Cobolt 06-MLD). The line illumination is formed with cylindrical lenses and the x-axis scanning was performed with a single-axis galvanometer scanner. The laser beam was reflected by a dichroic mirror and introduced into a Nikon Ti2-E inverted microscope. A prism-type spectrometer was built for the spectroscopic detection of fluorescence signals (prism: Thorlabs, PS855). The spectral detection range of the spectrometer was set to span from 410-622 nm. This range was achieved by using both a longpass filter (Semrock, LP02-407RU-25) and shortpass filters (Semrock, SP01-633RU-25). Here, the shortpass filter was used to block out near-infrared light used for the autofocus system of Ti2-E inverted microscope. The excitation beam was illuminated on the sample, and fluorescence signals were collected with an NA0.7 dry objective lens (Nikon, CFI Plan Fluor 60XC). The fluorescence signals were recorded with an EMCCD camera (Princeton Instruments, Pro-EM:1024). Image acquisition, image reconstruction, and image processing were performed by homebuilt software.

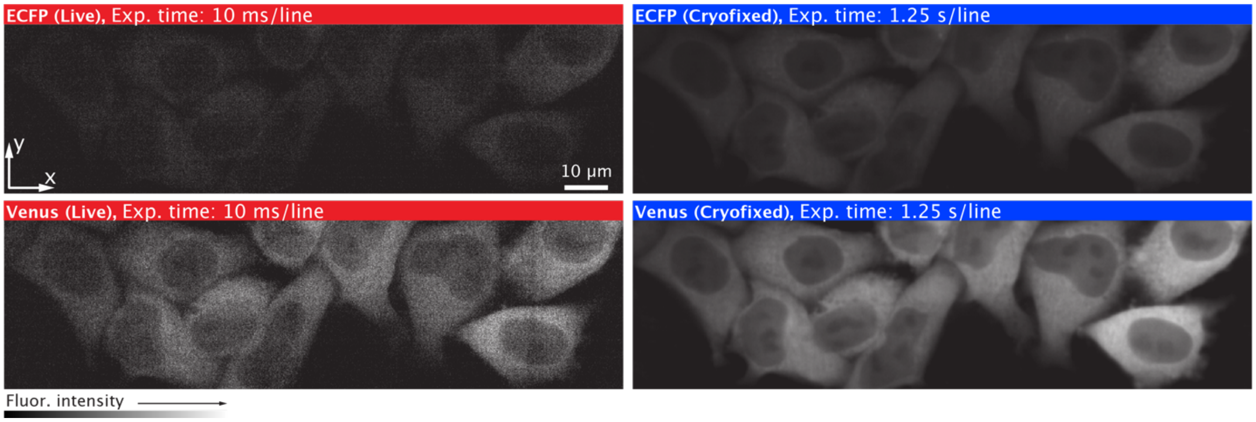


**Fig. S7 Fluorescence images of HeLa cells expressing YC3.60 before and after cryofixation.** The fluorescence intensity images were obtained using a hyperspectral slit-scanning fluorescence microscope shown in Fig. S6. The ratiometric fluorescence images in Fig. 3B of the main text were generated using these fluorescence intensity images. These fluorescence intensity images were reconstructed from measured fluorescence spectra and the background signals were subtracted in the manner described in Text S3h. Trehalose was not added to the buffer solution in this experiment.

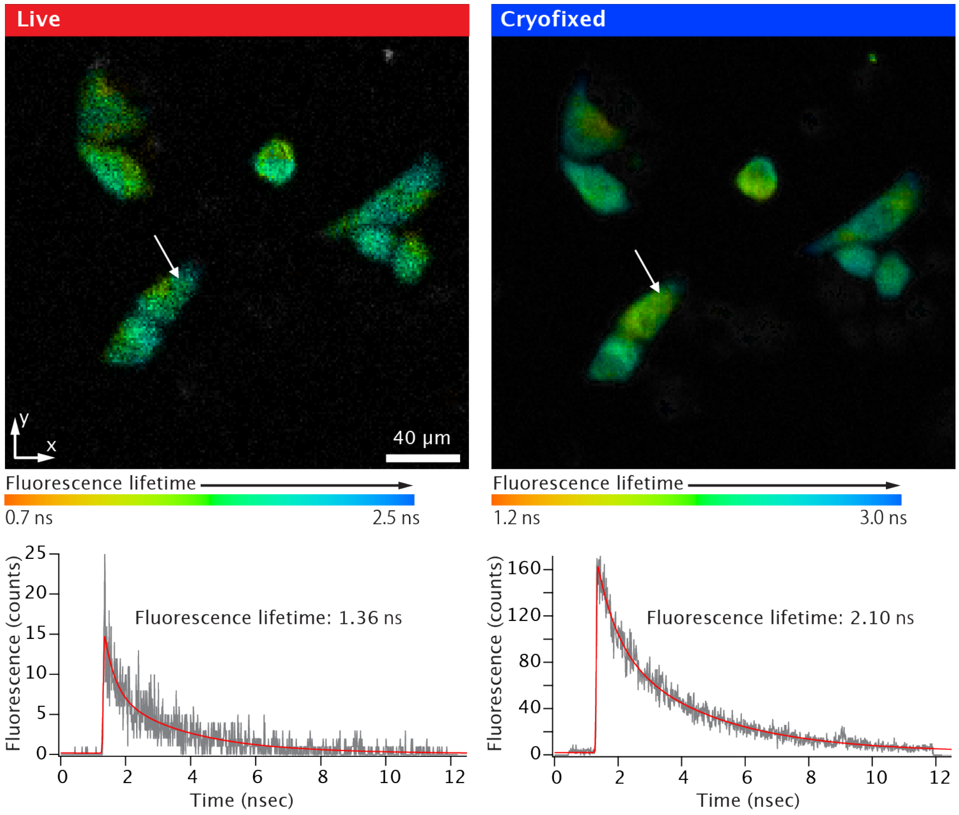


**Fig. S8.**

**Fluorescence lifetime images of HeLa cells expressing YC3.60 at room temperature and cryogenic conditions.** YC3.60 was excited with a femtosecond pulse laser at a center wavelength of 405 nm and fluorescence signals of ECFP were detected in a spectral band from 466-499 nm (Semrock, FF01-483/32-25). The pixel dwell times for 20 ℃ and -170 ℃ were 100 µs and 1 ms, respectively. After cryofixation, the fluorescence lifetime became 1.54 times longer. For this measurement, a conventional confocal laser-scanning fluorescence microscope was used. The laser source was an OPO system (Spectra Physics, Inspire HF100) seeded by an 80 MHz mode-locked Ti:sapphire laser (Spectra Physics, Mai-Tai). Laser scanning was performed with a 2-axis galvanometer scanner. The excitation laser beam was focused on a sample with an NA0.45 dry objective lens (Nikon, S Plan Fluor ELWD 20x) mounted in a Nikon Ti-E inverted microscope. Fluorescence signals were collected with the same objective lens and detected with a hybrid single-photon detector (Becker & Hickl, HPM-100). The detected fluorescence photons were counted using a time-correlated single-photon counting (TCSPC) board (Becker & Hickl, SPC-180NX). In our system, image acquisition was performed using SPCM data acquisition software (Becker &

Hickl), and data analysis was performed using SPCImage NG data analysis software (Becker & Hickl). Trehalose was not added to the buffer solution in this experiment.

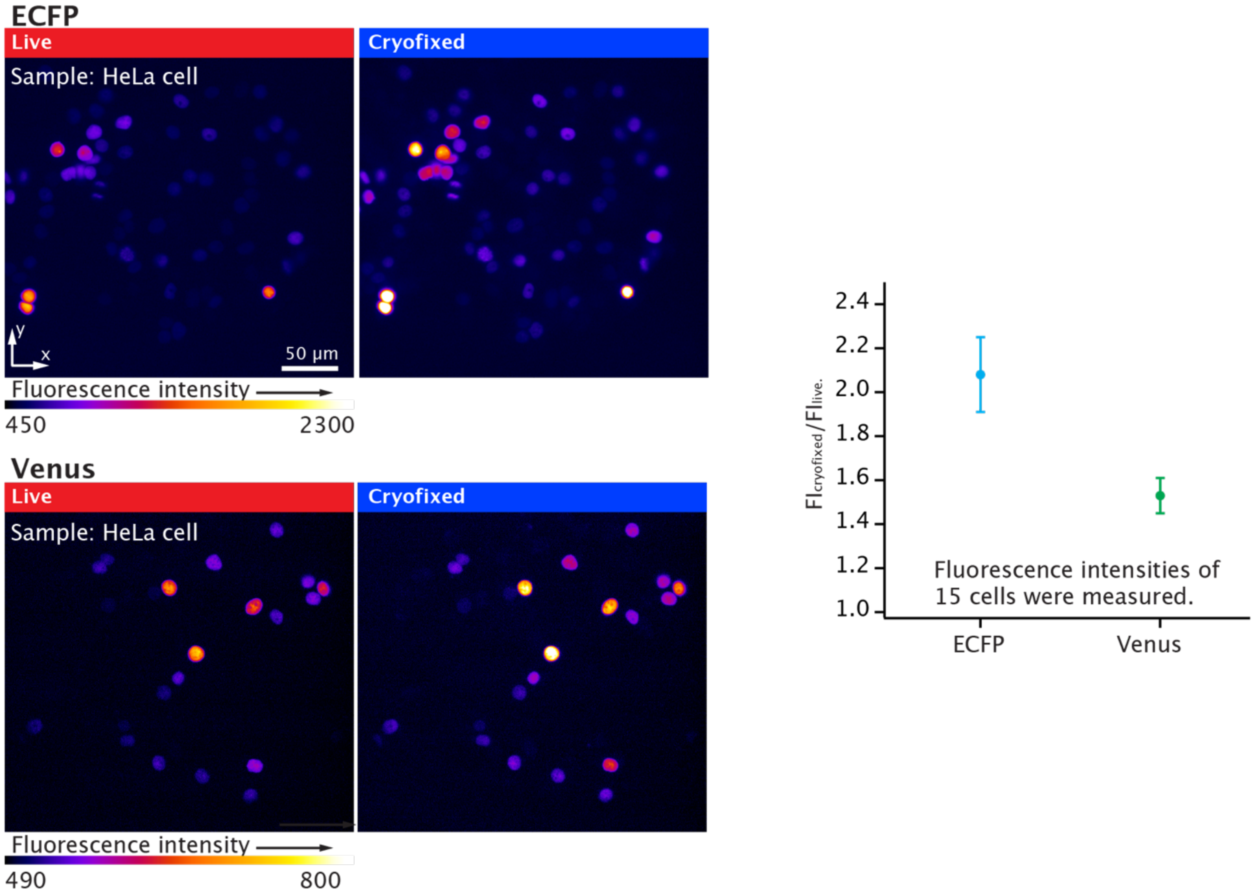


**Fig. S9.**

**Fluorescence images of HeLa cells expressing ECFP and Venus in nucleus before and after cryofixation.** The samples were observed with a conventional widefield fluorescence microscope equipped with a mercury lamp. From fluorescence intensities of 15 cells in each image, we confirmed that fluorescence intensities of ECFP and Venus were enhanced 2.08 and 1.53 times under the cryogenic condition, respectively. An NA0.45 dry objective lens (Nikon, S Plan Fluor ELWD 20x) was used for this observation. The excitation and detection wavelength bands for ECFP were 415-455 nm (Semrock, FF02-435/40-25) and 475-495 nm (Semrock, FF01-485/20-25), respectively. The excitation and detection wavelength bands for Venus were 464-499 nm (Semrock, FF01-482/35-25) and 516-556 nm (Semrock, FF01-536/40-25), respectively. Fluorescence signals were detected with an EMCCD camera (Andor, iXon Ultra 888) with a 100 ms exposure time. Trehalose was not added to the buffer solution in this experiment.

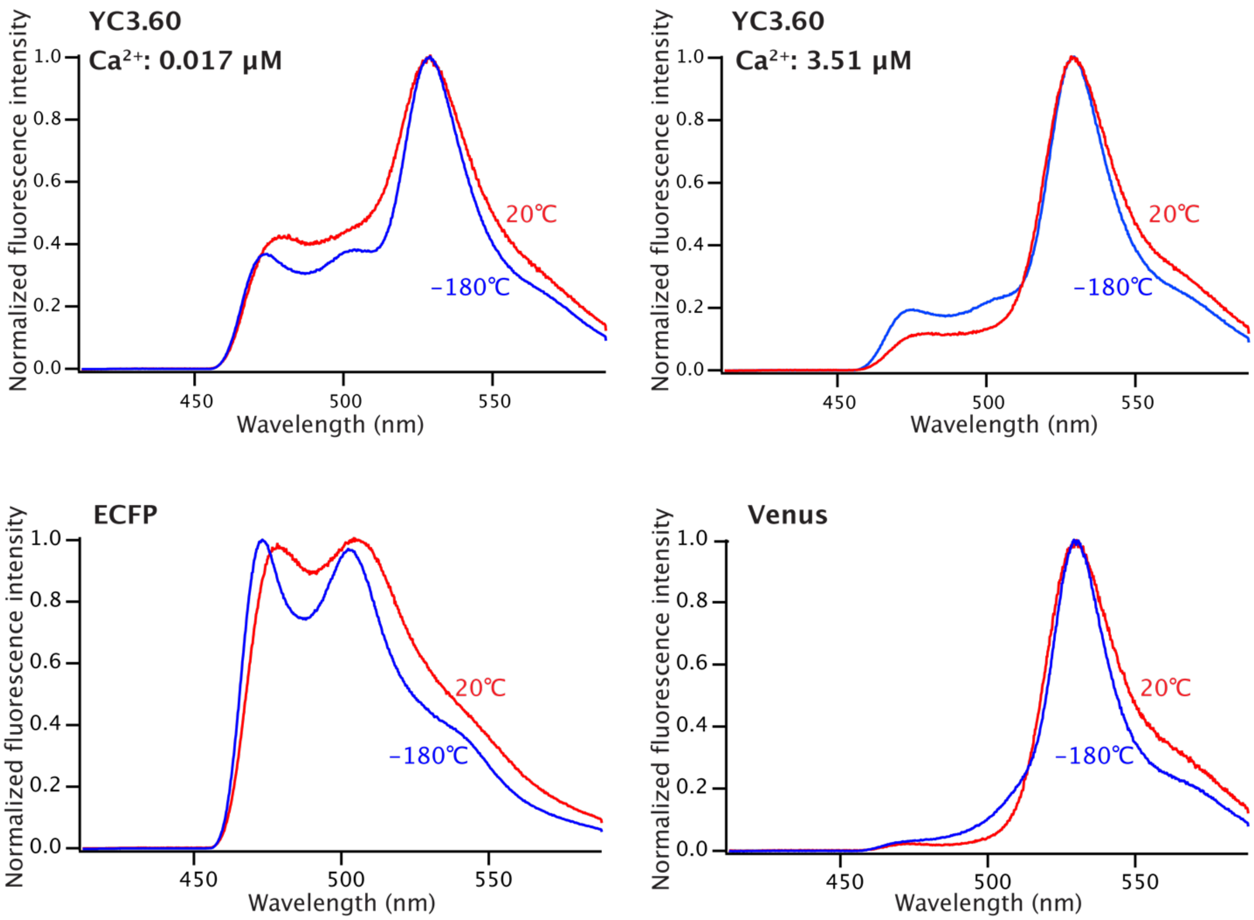


**Fig. S10.**

**Fluorescence spectra of YC3.60, ECFP, and Venus under before and after cryofixation.** Fluorescence spectra of YC3.60 at the $Ca^{2+}$ concentration of 0.017 and 3.51 μM were those obtained in the experiment shown in Fig. 3D. Fluorescence spectra of ECFP and Venus were observed with a conventional widefield fluorescence microscope equipped with a light emitting diode (LED) (Thorlabs, SOLIS-405C). The change in the fluorescence spectral shape of YC3.60 between room temperature and cryogenic conditions is considered to be attributed to the narrowing of the spectral peaks of ECFP and Venus under the cryogenic condition. Although the shape of the fluorescence emission spectrum of YC3.60 is slightly changed by cryofixation, the fluorescence signals can be detected by using the same set of optical filters as those used at room temperature. The excitation wavelength band was 400-410 nm (Semrock, FF01-405/10-25). The fluorescence signals were detected in a wavelength region above 461 nm (Semrock, LP03-458RU-25). An NA0.7 dry objective lens (Nikon, CFI Plan Fluor 60XC) was used for this observation.

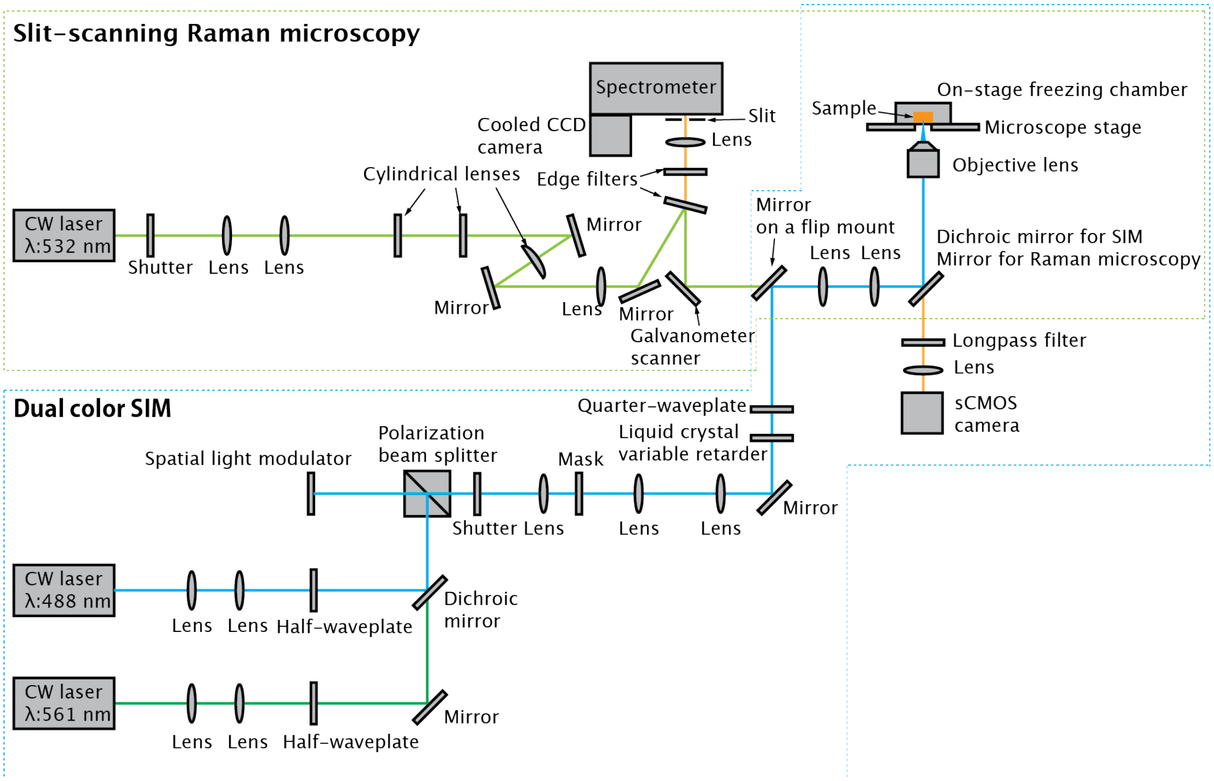


**Fig. S11.**

**Multimodal SIM/Raman microscope.** To perform multimodal SIM (fluorescence) and Raman imaging, an optical setup for a slit-scanning Raman microscopy is introduced into the SIM (Fig. S2). The optical configuration of the Raman microscope is similar to that previously reported in Ref. 47. The excitation wavelength for Raman imaging is 532 nm (Coherent, Verdi V18). The line illumination is formed by cylindrical lenses and the x-axis scanning is performed with a single-axis galvanometer scanner. In this setup, SIM (fluorescence) and Raman images are acquired separately by switching the excitation and detection optical paths by using a mirror mounted on a flip mount and rotating a filter cube turret equipped with filter cubes of the dichroic mirror for SIM and a mirror for Raman imaging. A slit is placed at the entrance of a spectrometer (Bunkoukeiki, MK-300), and the position of the slit is conjugated to the sample plane. Spontaneous Raman signals from line illumination on a sample are recorded with a cooled CCD camera (Princeton Instruments, PIXIS:400BR). Image acquisition, image reconstruction, and image processing such as background subtraction for Raman imaging were performed by the software we have developed. An NA0.95 dry objective lens (Nikon, CFI Plan Apo Lambda 60XC) was used for both the line illumination of excitation light and the collection of spontaneous Raman and fluorescence signals.

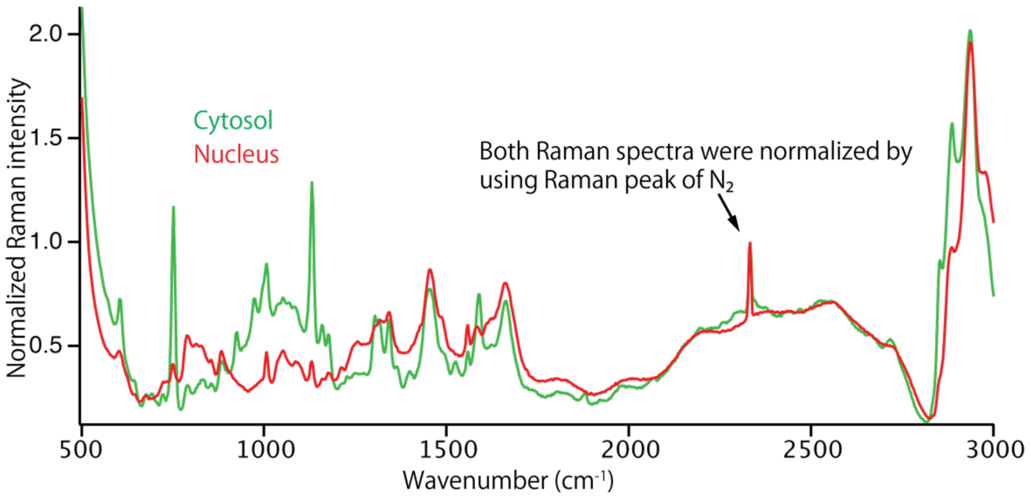


**Fig. S12.**

**Raman spectra of a HeLa cell under cryogenic condition.** These representative Raman spectra of cytosol (green) and nucleus (red) regions were obtained from a HeLa cell shown in Fig. 3E and were normalized by using the Raman peak of $N_2$ at 2332 $cm^{-1}$ indicated in the figure.

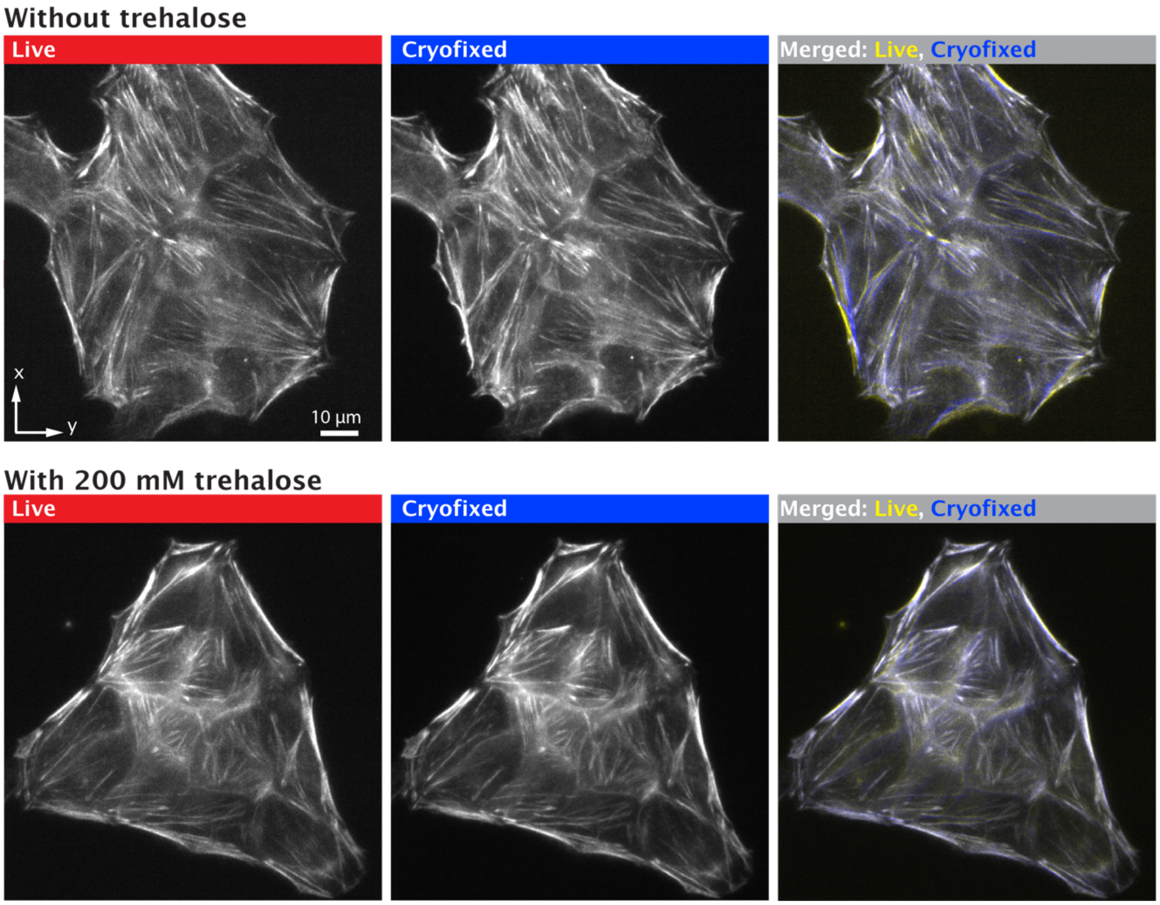


**Fig. S13.**

**Cryogenic fluorescence images of HeLa cells with and without cryoprotectant.** Actin filaments of HeLa cells labeled with SPY-555-actin (Spirochrome) were observed with a conventional widefield fluorescence microscopy equipped with a mercury lamp. Fluorescence images of actin filaments were observed with and without the presence of trehalose, a widely used cryoprotectant. As shown in the merged images, we observed slight differences of the cellular shapes between the fluorescence images before and after cryofixation and found that the alternation of cellular shapes was mitigated by adding trehalose to the buffer solution. Even with trehalose, the blurred parts exhibit the change in cell shapes. The blurred parts locate at the positions relatively far from the coverslip (in this observation, the focus position was set to be just above the coverslip), indicating that the parts surrounding the extracellular medium were affected more by freezing. This also implies that the difference in volume changes of the cell body and the extracellular solution caused the slight change in the morphology.

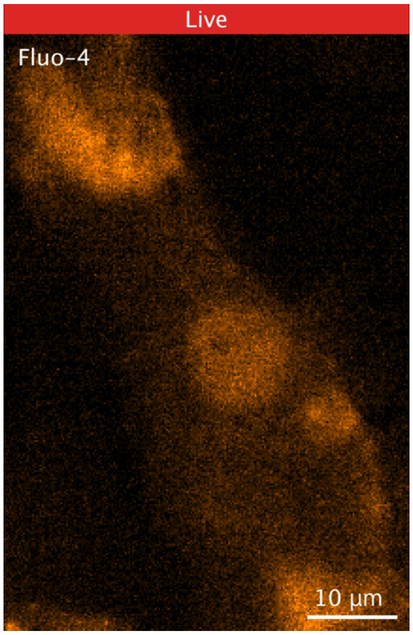


**Movie S1.**
**Cryofixation of $Ca^{2+}$ wave in neonatal rat cardiomyocytes (Fig. 1C).**

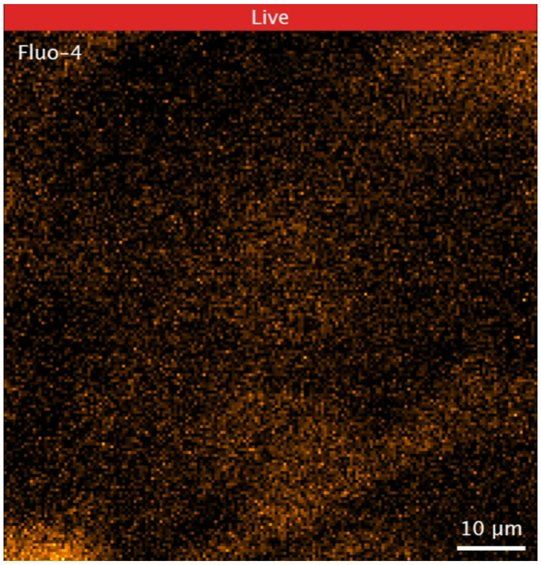


**Movie S2.**
**Cryofixation of $Ca^{2+}$ wave in neonatal rat cardiomyocytes without cryoprotectant (Fig. 1G).**

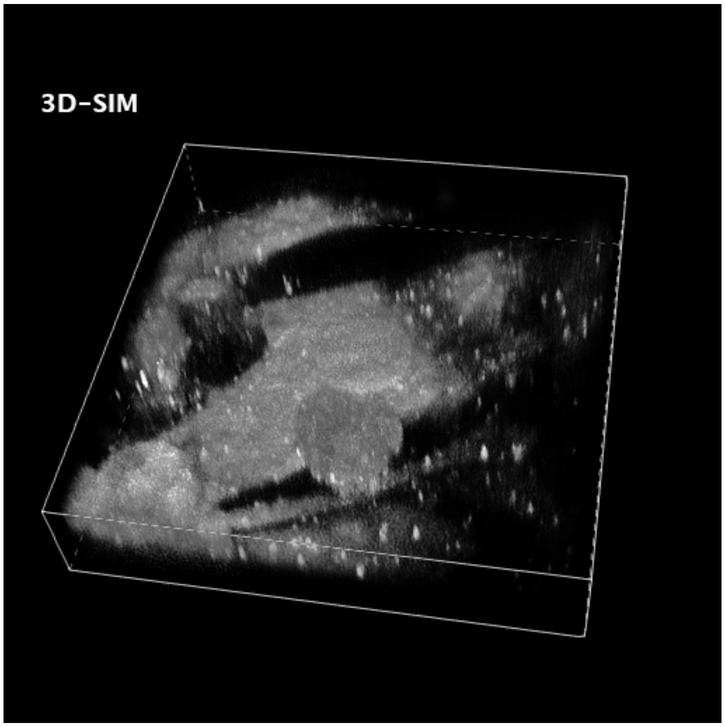


**Movie S3.**

**3D visualization of cryofixed $Ca^{2+}$ waves in neonatal rat cardiomyocytes (Fig. 1G)**

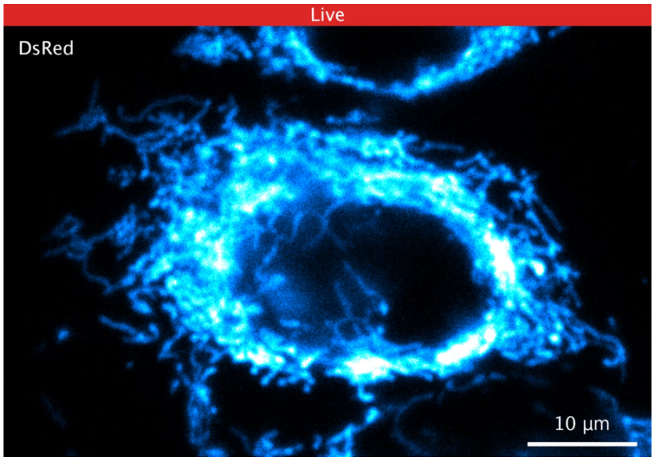


**Movie S4.**
**Cryofixation of mitochondria in HeLa cells without cryoprotectant (Fig. 1H).**

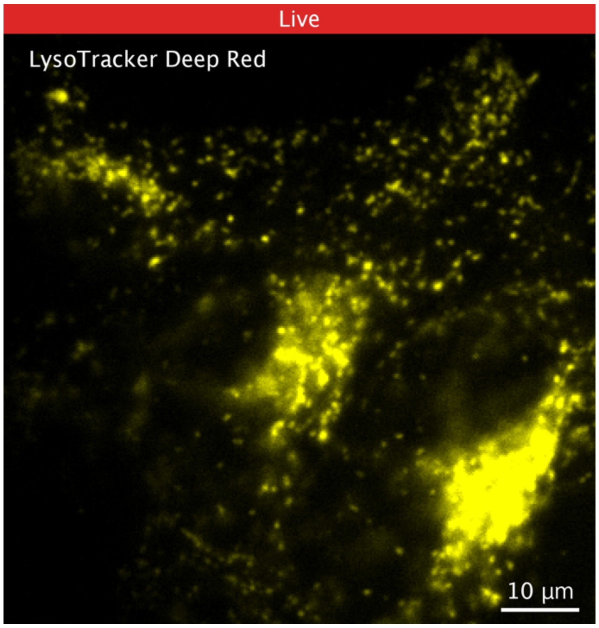


**Movie S5.**

**Cryofixation of lysosomes in HeLa cells without cryoprotectant.** HeLa cells labeled with LysoTracker Deep Red (Thermo Fisher Scientific, L12492) were observed using a conventional inverted wide-field fluorescence microscope equipped with a mercury lamp. As shown in this movie, the dynamic motion of the lysotrackers was observed at room temperature and was immediately stopped by cryofixation. The excitation and detection wavelength bands were 609/54 nm (Semrock, FF01-609/54-25) and 708/75 nm (Semrock, FF01-708/75-25). The sample was observed with an NA0.7 dry objective lens (Nikon, CFI Plan Fluor 60XC 60x), and fluorescence images were acquired with an sCMOS camera (Hamamatsu Photonics, ORCA Flash4.0 V3), with an exposure time of 100 ms. The fluorescence intensities of each image were normalized for visualization in the same manner as shown in Figure 1C.

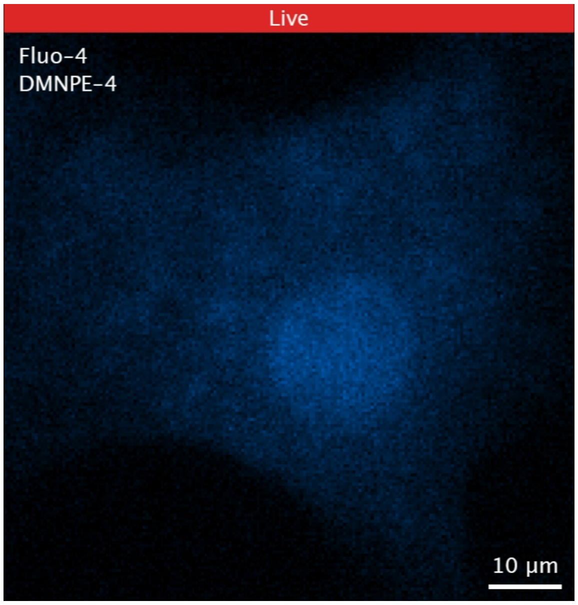


**Movie S6.**

**Time-deterministic cryofixation of $Ca^{2+}$ wave induced by uncaging $Ca^{2+}$ from a caged $Ca^{2+}$ compound with UV light irradiation (Fig. 2A).**

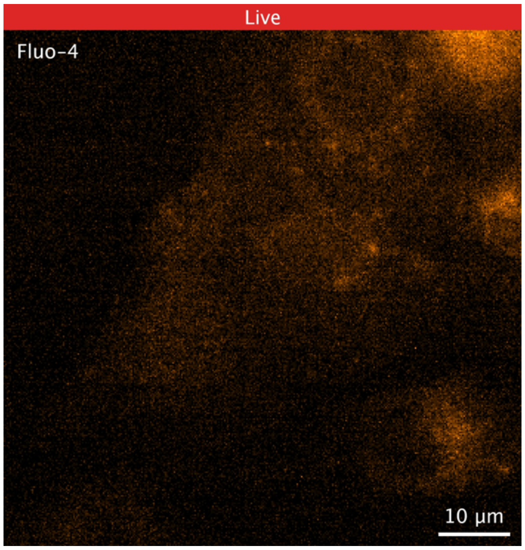


**Movie S7.**

**Time-deterministic cryofixation of neonatal rat cardiomyocytes at the contraction phase (Fig. 2B).**

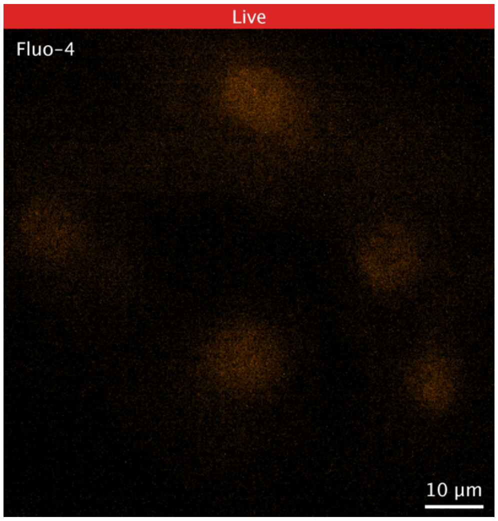


**Movie S8.**

**Time-deterministic cryofixation of neonatal rat cardiomyocytes at the relaxation phase (Fig. 2C).**
